# scAMACE: Model-based approach to the joint analysis of single-cell data on chromatin accessibility, gene expression and methylation

**DOI:** 10.1101/2021.03.29.437485

**Authors:** Jiaxuan Wangwu, Zexuan Sun, Zhixiang Lin

**Affiliations:** Department of Statistics, The Chinese University of Hong Kong, Hong Kong SAR, China

**Keywords:** Bayesian statistics, Clustering, Data integration, Multi-omics, Single-cell genomic

## Abstract

The advancement in technologies and the growth of available single-cell datasets motivate integrative analysis of multiple single-cell genomic datasets. Integrative analysis of multimodal single-cell datasets combines complementary information offered by single-omic datasets and can offer deeper insights on complex biological process. Clustering methods that identify the unknown cell types are among the first few steps in the analysis of single-cell datasets, and they are important for downstream analysis built upon the identified cell types. We propose scAMACE for the integrative analysis and clustering of single-cell data on chromatin accessibility, gene expression and methylation. We demonstrate that cell types are better identified and characterized through analyzing the three data types jointly. We develop an efficient expectation-maximization (EM) algorithm to perform statistical inference, and evaluate our methods on both simulation study and real data applications. We also provide the GPU implementation of scAMACE, making it scalable to large datasets. The software and datasets are available at https://github.com/cuhklinlab/scAMACE_py (python implementation) and https://github.com/cuhklinlab/scAMACE (R implementation).

## 1 Introduction

Recent developments in single-cell technologies enable multiple measurements of different genomic features (Lahnemann *et al*., 2020). Sequencing technologies include single-cell RNA sequencing (scRNA-seq) which measures transcription, single-cell ATAC sequencing (scATAC-seq) and the assay based on combinatorial indexing (sci-ATAC-seq) (Cusanovich *et al*., 2018b) that measure chromatin accessibility, and single-nucleus methylcytosine sequencing (snmC-seq) (Luo *et al*., 2017) which meansures methylome at the single cell resolution. High technical variation is presented in single-cell datasets due to the limited amount of genomic materials and the experimental procedures to amplify the signals (Lahnemann *et al*., 2020).

Because cell types are usually unknown beforehand, clustering methods are needed to identify the cell types. Majority of existing clustering algorithms only take one single dataset as input. Beside the widely used K-Means clustering algorithm, hierarchical clustering (Ward, 1963) forms hierarchical groups of mutually exclusive subsets on the basis of their similarity with respect to specified characteristics by considering the union of all possible 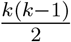 pairs and accepting the union with which an optimal value of the objective function is associated. Spectral Clustering (Ng *et al*., 2001) uses the top *k* eigenvectors of a matrix derived from the distance between points simultaneously for clustering. Several algorithms are developed specifically for scRNA-seq data. SC3 (Kiselev *et al*., 2017) combines multiple clustering outcomes through a consensus approach. SIMLR (Wang *et al*., 2017) learns a distance metric by multiple kernels and clusters with affinity propagation. CIDR (Lin *et al*., 2017) imputes the gene expression profiles, calculates the dissimilarity based on the imputed gene expression profiles for every pair of single cells, performs principal coordinate analysis using the dissimilarity matrix, and finally performs clustering using the first few principal coordinates. SOUP (Zhu *et al*., 2019) semi-softly classifies both pure and intermediate cell types: it first identifies the set of pure cells by special block structure and estimates a membership matrix, then estimates soft membership for the other cells. For the analysis of single-cell chromatin accessibility data, scABC (Zamanighomi *et al*., 2018) first weights cells and applies weighted K-medoids clustering, then calculate landmarks for each cluster, and finally clusters the cells by assignment to the closest landmark based on Spearman correlation. Cusanovich (Cusanovich *et al*., 2018a) makes use of singular value decomposition on TF-IDF transformed matrix and density peak clustering algorithm. cisTopic (Bravo Gonzá lez-Blas *et al*., 2019) uses latent Dirichlet allocation with a collapsed Gibbs sampler to iteratively optimize the region-topic distribution and the topic-cell distribution. SCALE (Xiong *et al*., 2019) combines the variational autoencoder framework with the Gaussian Mixture Model which extracts latent features that characterize the distributions of input scATAC-seq data, and then uses the latent features to cluster cell mixtures into subpopulations. Clustering methods are also developed for single-cell methylation data. BPRMeth (Kapourani and Sanguinetti, 2016) uses probabilistic machine learning to extract higher order features across a defined region and to cluster promoter-proximal regions by Binomial distributed probit regression (BPR) and mixture modeling. PDclust (Hui *et al*., 2018) leverages the methylation state of individual CpGs to obtain pairwise dissimilarity (PD) values, and calculates Euclidean distances between each pair of cells using their PD values and performed hierarchical clustering. Melissa (Kapourani and Sanguinetti, 2019) implements a Bayesian hierarchical model that jointly learns the methylation profiles of genomic regions of interest and clusters cells based on their genome-wide methylation patterns. pCSM (Yin *et al*., 2019) implements a semi-reference-free procedure to perform virtual methylome dissection using the nonnegative matrix factorization algorithm. It first determines putative cell-type-specific methylated loci and then clusters the loci into groups based on their correlations in methylation profiles.

Studies based on single-omic data provide only a partial landscape of the entire cellular heterogeneity (Ma *et al*., 2020). High technical noise and the growth of available datasets measuring different genomic features encourage integrative analysis (Lahnemann *et al*., 2020). By combining complementary information from multiple datasets, the cell types may be better seperated and characterized (Corces *et al*., 2016; Duren *et al*., 2017). The integrative analysis of gene expression and chromatin activity may better define cell types and lineages, especially in complex tissues (Duren *et al*., 2018). Seurat V3 (Stuart *et al*., 2019) uses Canonical Correlation Analysis (CCA) to reduce the dimension of the datasets. It identifies the pairwise correspondences of single cells across datasets, termed ‘anchors’, and then transfers labels from a reference dataset onto a query dataset. coupleNMF (Duren *et al*., 2018) is based on the coupling of two non-negative matrix factorizations, where a ‘soft’ clustering can be obtained following the matrix factorizations. It enables integrative analysis of scRNA-seq and scATAC-seq data. LIGER (Welch *et al*., 2019) integrates multimodal datasets via integrative non-negative matrix factorization (iNMF) to learn a low-dimensional space defined by dataset-specific factors and shared factors across datasets, and then build a neighborhood graph based on the shared factors to identify joint clusters by performing community detection on this graph. scACE (Lin *et al*., 2020) is a model-based approach that jointly analyzes single-cell chromatin accessibility and scRNA-Seq data, and it quantifies the uncertainty of cluster assignments. MAESTRO (Wang *et al*., 2020) integrates scRNA-seq and scATAC-seq data from multiple platforms. It also provides comprehensive functions for pre-processing, alignment, quality control, and quantification of expression and accessibility. coupleCoC (Zeng *et al*., 2020) performs co-clustering of the cells and the features simultaneously in the source data and the target data, and it also matches the cell clusters between the source data and the target data through minimizing the distribution divergence. scMC (Zhang and Nie, 2021) integrates multiple scRNA-Seq datasets or multiple scATAC-Seq datasets, where it learns biological variation via variance analysis to subtract technical variation inferred in an unsupervised manner. The three data types, including gene expression, chromatin accessibility and methylation, have distinct characteristics and complex relationships with each other. The aforementioned methods for integrative analysis are not designed to integrate all three data types. Moreover, these methods (except scACE) do not provide statistical inference on the cluster assignments, which may be important when there are cells at the intermediate stages during development.

In this work, we extend scACE (Lin *et al*., 2020) to scAMACE (integrative Analysis of single-cell Methylation, chromatin ACcessibility, and gene Expression). scAMACE considers the biological and technical variabilities when integrating multiple data types, and it can provide statistical inference on the assignment of clusters. We reason that by combining complementary biological information from multiple data types, better cell type seperation can be achieved. We present our model in Section 2, and statistical inference using the Expectation-Maximization (EM) algorithm in Section 3. Simulation study and real data applications are presented in Sections 4 and 5, respectively. The conclusion is presented in Section 6.

## 2 Methods

An overview of scAMACE is presented in Fig. 1.

**Figure 1:**
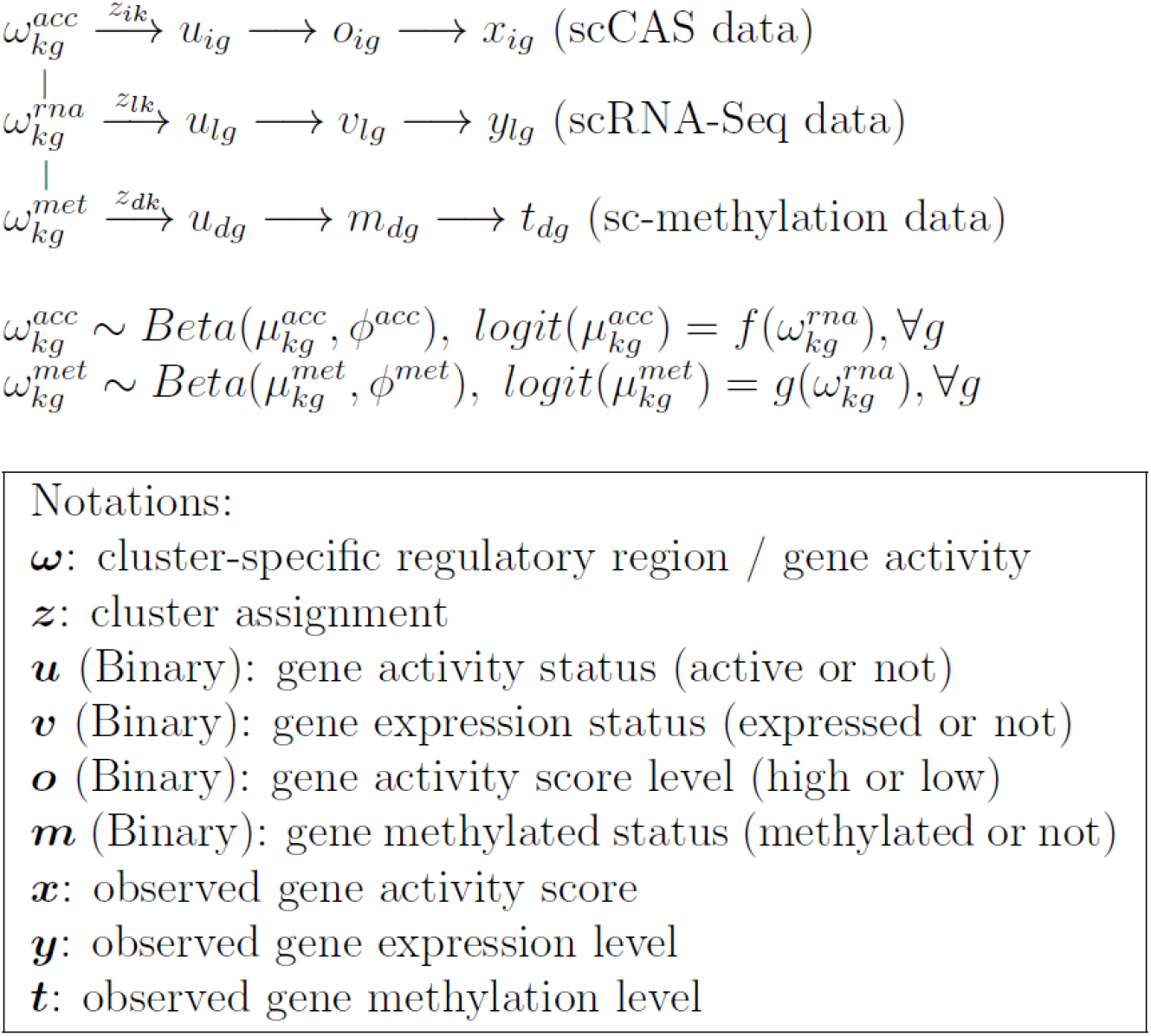
Graphical representation of scAMACE.

### 2.1 Model for scRNA-Seq data

The model specification for scRNA-Seq data is as the following.

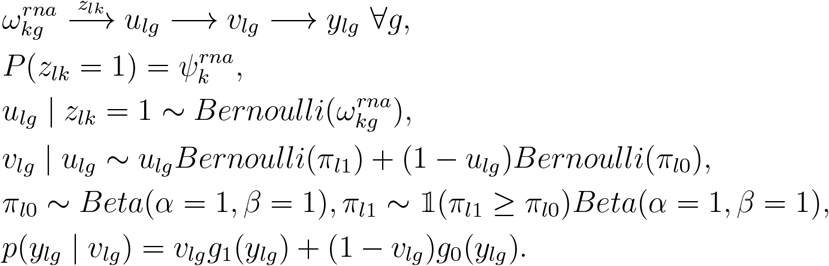

We assume that there are *K* cell clusters in total, the random variable *z*_*lk*_ denotes whether cell *l* belongs to cluster *k* ∈ {1, …, *K*}, and ***z***_*l·*_ follows categorical distribution with probability 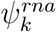 for cluster *k*.

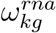 denotes the probability that gene *g* is active in cluster *k. u*_*lg*_ is a binary latent variable representing whether gene *g* is active in cell *l* and *u*_*lg*_ = 1 represents that it is active. *v*_*lg*_ denotes whether gene *g* is expressed in cell *l* and *v*_*lg*_ = 1 represents that it is expressed.

When gene *g* is active in cell *l* (*u*_*lg*_ = 1), the probability that gene *g* is expressed in cell *l* (*v*_*lg*_ = 1) is *π*_*l*1_, while the probability that gene *g* is expressed is *π*_*l*0_ if the gene is not active (*u*_*lg*_ = 0). Since genes are more likely to be expressed when they are active, we assume that *π*_*l*1_ ≥ *π*_*l*0_ and the prior distributions of *π*_*l*1_ and *π*_*l*0_ are assumed to be flat.

Let *y*_*lg*_ denote the observed gene expression for gene *g* in cell *l* (after normalization to account for sequencing depth and gene length), and we assume that *y*_*lg*_ | *v*_*lg*_ follows a mixture distribution, where *g*_1_(.) and *g*_0_(.) are density functions of the expression level conditional on *v*_*lg*_.

### 2.2 Model for single-cell chromatin accessibility (scCAS) data

The model specification for scCAS data is as the following.

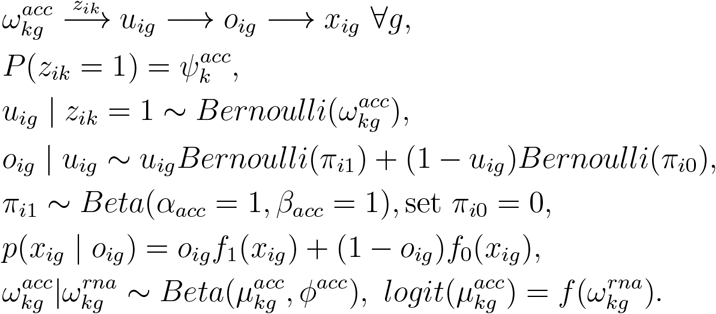

The random variables 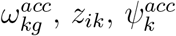 and *u*_*ig*_ have similar interpretations to their corresponding variables in the model for scRNA-Seq data. We use a different notation *i* to represent that the cells in the scCAS data are different from the cells in the scRNA-Seq data.

*x*_*ig*_ denotes the observed gene score for gene *g* in cell *i*. The gene score summarizes the accessibility of the regions around the gene body (Cusanovich *et al*., 2018a). We model it by a mixture distribution with density functions *f*_1_(.), *f*_0_(.), and binary latent variable *o*_*ig*_. *o*_*ig*_ = 1, and 0 represent the mixture components with high (*f*_1_) and low (*f*_0_) gene scores, respectively. Accessibility tends to be positively associated with activity of the gene. We model this positive relationship by the distribution *o*_*ig*_|*u*_*ig*_. When gene *g* is active in cell *i* (*u*_*ig*_ = 1), the probability that it has high gene score (*o*_*ig*_ = 1) is *π*_*i*1_; When gene *g* is inactive in cell i (*u*_*ig*_ = 0), the probability that it has high gene score (*o*_*ig*_ = 1) is *π*_*i*0_. We assume that *π*_*i*1_ ≥ *π*_*i*0_ to represent the positive relationship. In practice, we found that fixing *π*_*i*0_ = 0 leads to good real data performance, and we set *π*_*i*0_ = 0 by default. The prior distribution *π*_*i*1_ ∼ *Beta*(*α* = 1, *β* = 1). In real data example 1, the observed data is promoter accessibility and we use the same model as that for gene score.

We assume that 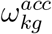 follows Beta distribution with mean 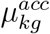 and precision *ϕ*^*acc*^. The variable 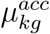 is connected with 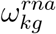 in scRNA-Seq data through the logit function: 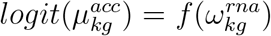. Details on the specification of *f* (·) are presented in Section 2.6.

### 2.3 Model for single-cell methylation data

The model specification for sc-methylation data is as the following.

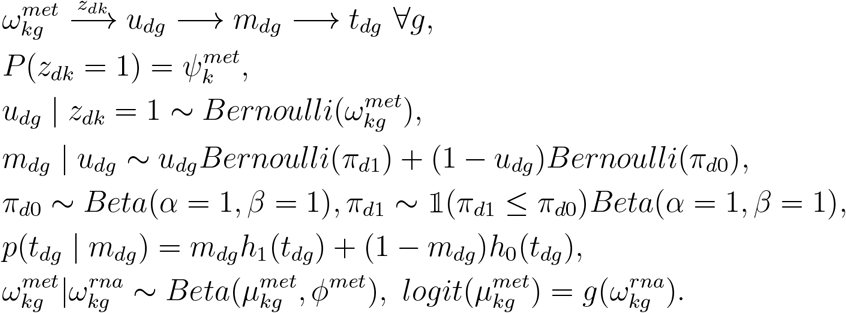

The random variables 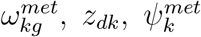, and *u*_*dg*_ have similar interpretations to their corresponding variables in the model for scRNA-Seq data. We use a different notation *d* to represent that the cells in the sc-methylation data are different from the cells in the scRNA-Seq data.

The binary random variable *m*_*dg*_ denotes whether gene *g* is methylated in cell *d*, and *m*_*dg*_ = 1 represents that it is methylated. Methylation of a gene (promoter methylation/gene body methylation) tends to be negatively associated with activity of the gene, and we model this negative relationship with the model *m*_*dg*_ | *u*_*dg*_: when the gene *g* is active in cell *d* (*u*_*dg*_ = 1), it is less likely to be methylated (*m*_*dg*_ = 1), as we assume that *π*_*d*1_ ≤ *π*_*d*0_.

*t*_*dg*_ denotes the observed methylation level for gene *g* in cell *d*, and we assume that *t*_*dg*_ | *m*_*dg*_ follows a mixture distribution, where *h*_1_(.) and *h*_0_(.) are density functions conditional on *m*_*dg*_. The technologies/features differ for the two real data applications to be presented: promoter methylation for the gene (Pott, 2017), and gene body methylation at non-CG sites (Luo *et al*., 2017).

Similar to scCAS data, we connect 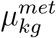, which is the mean of 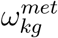, and 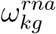 through the logit function: 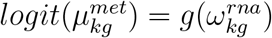. Details on specification of *g*(·) are presented in Section 2.6.

### 2.4 More on model specification

Methylation and chromatin accessibility regulate gene expression biologically. Our model is specified in the reverse order, so gene expression plays a central role. This is because scRNA-Seq data is usually less noisy compared with scCAS data and sc-methylation data, the model specified this way will improve the clustering performance of scCAS data and sc-methylation data, without sacrificing much the clustering performance of scRNA-Seq data.

### 2.5 Prior specifications

We assume the following priors for 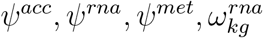.

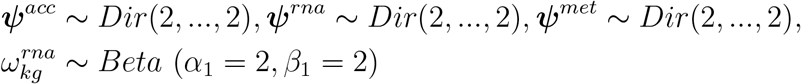

The prior specification *Beta* (*α* = 2, *β* = 2) improves the stability of the EM algorithm in Section 3 over uniform distritbution.

### 2.6 Determination of 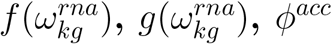 and *ϕ*^*met*^

We assume that 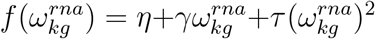 and 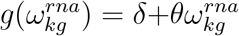. The parameters {*η, γ, τ, δ, θ, ϕ*^*acc*^, *ϕ*^*met*^} are estimated empirically from the datasets. We first set the number of clusters *K* = 1 and use the model to estimate 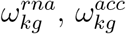 and 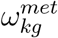 separately without considering the links on ***ω*** across the three datasets, and then fix 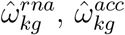 and 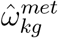 to estimate {*η, γ, τ, δ, θ, ϕ*^*acc*^, *ϕ*^*met*^} by beta regression (Silvia and Francisco, 2004). The rationale for fixing *K* = 1 to estimating the parameters in the functions *f* (.) and *g*(.) that link the three modalities is that the majority of the features may not change much across the cell types. We fix 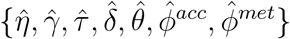 when implementing the EM algorithm in Section 3. Estimating {*η, γ, τ, δ, θ, ϕ*^*acc*^, *ϕ*^*met*^} separately from the EM algorithm improves computational efficiency and avoids problematic local modes. Distributions of 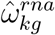 v.s. 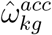 and 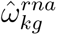 v.s. 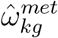 for the two real data applications are presented in Supplementary Materials Figures S.4 and S.5, we can see from Figures S.4 and S.5 that the linear and quadratic models capture the trends on how 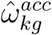 and 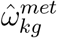 changes with 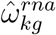.

### 2.7 The mixture components

For scCAS data, we apply *f*_1_(*x*) = 0, *f*_0_(*x*) = 1 if *x* = 0 and *f*_1_(*x*) = 1, *f*_0_(*x*) = 0 if *x >* 0, due to the sparsity of the data matrix.

For scRNA-Seq data, we first normalize read counts to TPM (transcripts per million) or FPKM (fragments per kilobase of exon model per million reads mapped) to account for sequencing depth and gene length, then fit a two-component gamma mixture model for the nonzero entries, through pooling ln(TPM+1) or ln(FPKM+1) over all the samples, and then the remaining zero entries are merged with the mixture component that has a smaller mean. The log transformation takes into account the very large values in the data matrix.

sc-methylation data represents the proportion of methylated sites within a given genomic interval, where the entries in the data matrix take values between 0 and 1. In the two real data applications, majority of entries in the data matrix take small values, for the methylation data in each cell, we first divide the entries by (1 — entries) to map them into [0, ∞). We then normalize the entries by dividing the median of non-zero entries in each cell, and then take square of the entries to boost the signals. This transformation helps to align the three modalities and it improves the clustering results significantly (Supplementary Materials Tables S.5 and S.6). Because the transformed entries represent the relative evidence of the methylation status, we input the transformed entries directly as the ratio 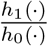 in the EM algorithm. Histograms for the distributions of the sc-methylation data are presented in the Supplementary Materials Figure S.1.

### 2.8 Feature selection

scRNA-Seq data is usually the least noisy data type, compared with scCAS and sc-methylation data. We use scRNA-Seq data for feature selection before implementing scAMACE. We first cluster scRNA-Seq data with SC3 and then use the cluster assignments to select top 1,000 features with large mean shift across different clusters. More specifically, denote the data matrix as ***X***_*n×p*_ (*x*_*ij*_ denotes the observation for the *i*-th cell and *j*-th feature), the cluster assignments as ***L***_*n×*1_ (*l*_*i*_ = *k* denotes that the *i*-th cell belongs to the *k*-th cluster) and total number of clusters as *K*. For feature *j*, we first calculate the difference between the mean of the cells within one cell type and the mean of cells in other cell types; the differences are represented as ***D***(*j*) = (*d*_1*j*_, …, *d*_*Kj*_), where 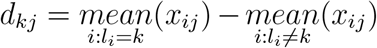. We take the maximum entry in *D*(*j*): *m*(*j*) = max_*k*_ ***D***(*j*). When *m*(*j*) is large, it represents that feature *j* has high expression in one cluster, compared with all other clusters. Finally, we select the top 1,000 features with highest values in *m*(*j*).

### 2.9 Determination of the number of clusters *K*

We determine the number of clusters *K* for the three single-cell datasets seperately before we apply scAMACE. We first run K-Means for each *K* and calculate the average silhouette width of observations (Kaufman and Rousseeuw, 1990). Silhouette width measures how well an observation has been classified. For each observation *i*, the silhouette value *s*(*i*) is calculated as follows. First denote by *A* the cluster to which observation *i* has been assigned and then calculate

*a*(*i*) = average Euclidean distance of *i* to all other objects of *A*.

Now consider any cluster *C* different from *A* and define

*d*(*i, C*) = average Euclidean distance of *i* to all objects of *C*.

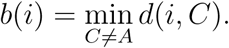

Then 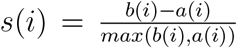. When cluster *A* contains only a single observation, we simply set *s*(*i*) = 0. The average of *s*(*i*) for *i* = 1, 2, …, *n* is denoted by 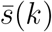, and it is called the average silhouette width for the entire data set. 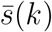 is used for the selection of *K*. Higher value in 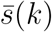 indicates better clustering outcome. We select *K* that has the maximum average silhouette width. Details for selecting *K* in the two real data applications are presented in Supplementary Materials Figures S.2 and S.3. When the similarity of the cell types is high, the Silhoutte method may choose a smaller *K* than the number of cell types (Figure S.3), and we may choose a larger *K* instead.

## 3 Statistical inference: EM algorithm

Given the observed scCAS data ***X***, scRNA-Seq data ***Y***, and sc-methylation data ***T***, we treat the latent variables **Γ** = {***Z, U***, ***O, V***, ***M***} as missing data, and use the Expectation-Maximization (EM) algorithm to estimate the parameters **Φ**={***ψ***^*acc*^, ***ω***^*acc*^,***π***_*i*_,***ψ***^*rna*^,***ω***^*rna*^, ***π***_*l*_, ***ψ***^*met*^, ***ω***^*met*^,***π***_*d*_}. The Q-function is ***Q***(**Φ**|**Φ**_*old*_) = 𝔼_*old*_(*ln*(***P*** (**Φ, Γ**|*obs*.))), where the expectation is over **Γ** under distribution ***P*** (**Γ**|**Φ**^*old*^, *obs*.).

In the M-step, we maximize ***Q***(**Φ**|**Φ**_*old*_) with respect to **Φ** and update parameters as follows.

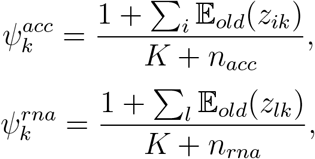

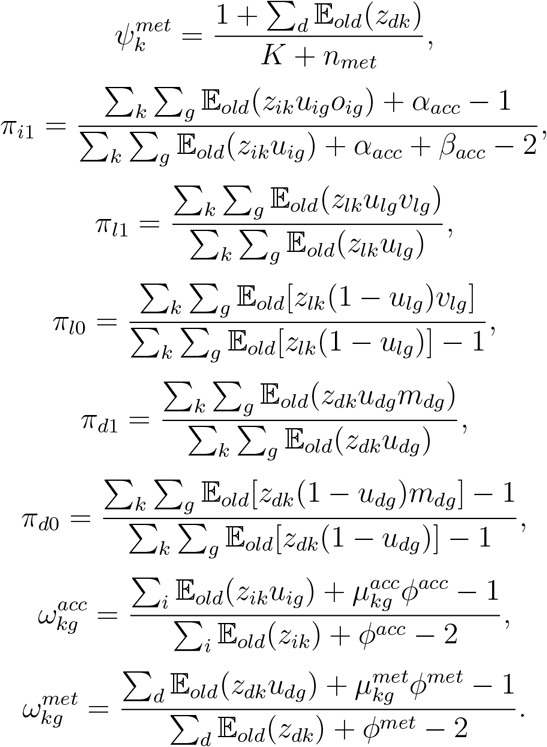

We use grid search to update 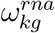 because its optimal value does not have an explicit form.

We iterate between E-step and M-step until converge. 𝔼 (***Z***_*i*._), 𝔼 (***Z***_*l*._) and 𝔼 (***Z***_*d*._) in the last iteration are used for clustering. Details for the derivations are presented in the Supplementary Materials.

## 4 Simulation studies

To validate scAMACE, we generated three different types of simulated data ***x, y*** and ***t*** following the model assumption. In the simulated data, the sample sizes *n*_*x*_ = 900, *n*_*y*_ = 1100, and *n*_*t*_ = 1000. The number of features *p* = 1000. The number of clusters *K*_*x*_ = *K*_*y*_ = *K*_*t*_ = 3, and 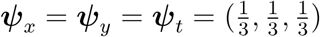.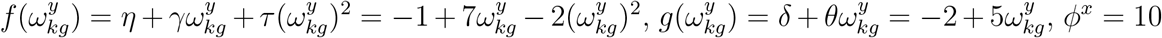, and *ϕ*^*t*^ = 10. The detailed simulation scheme is presented in the Supplementary Materials.

For the first data type, x, we set *f*_1_(*x*) = 0 if *x* = 0, and *f*_0_(*x*) = 0 if *x* = 1. We fit a two-component gamma mixture model for ***y*** using ‘gammamixEM’ in R (Young *et al*., 2019) and beta mixture model for ***t*** using ‘betamix’ in R (Cribari-Neto and Zeileis, 2010; Grun *et al*., 2012) to estimate the mixture densities. We apply the method in Section 2.6 to estimate parameters in 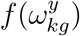 and 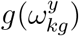. We then implement scAMACE using the estimated densities and 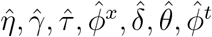.

We use purity, rand index, adjusted rand index and normalized mutual information to evaluate the clustering results. We implement scAMACE either on the three data types seperately (‘scAMACE (seperate)’) without borrowing information or jointly (‘scAMACE (joint)’). Table 1 presents the simulation results. We also compared scAMACE with other exsiting methods under four additional simulation schemes: imbalanced datasets where the number of cells varies across the three datasets (Supplementary Materials Table S.1), unequal numbers of clusters in the three datasets (Supplementary Materials Table S.2), imbalanced cluster sizes (Supplementary Materials Table S.3) and smaller number of features (Supplementary Materials Table S.4). scAMACE performs the best compared with the other methods in the above simulation settings. This is likely due to integration of information from all three data sets.

**Table 1:**
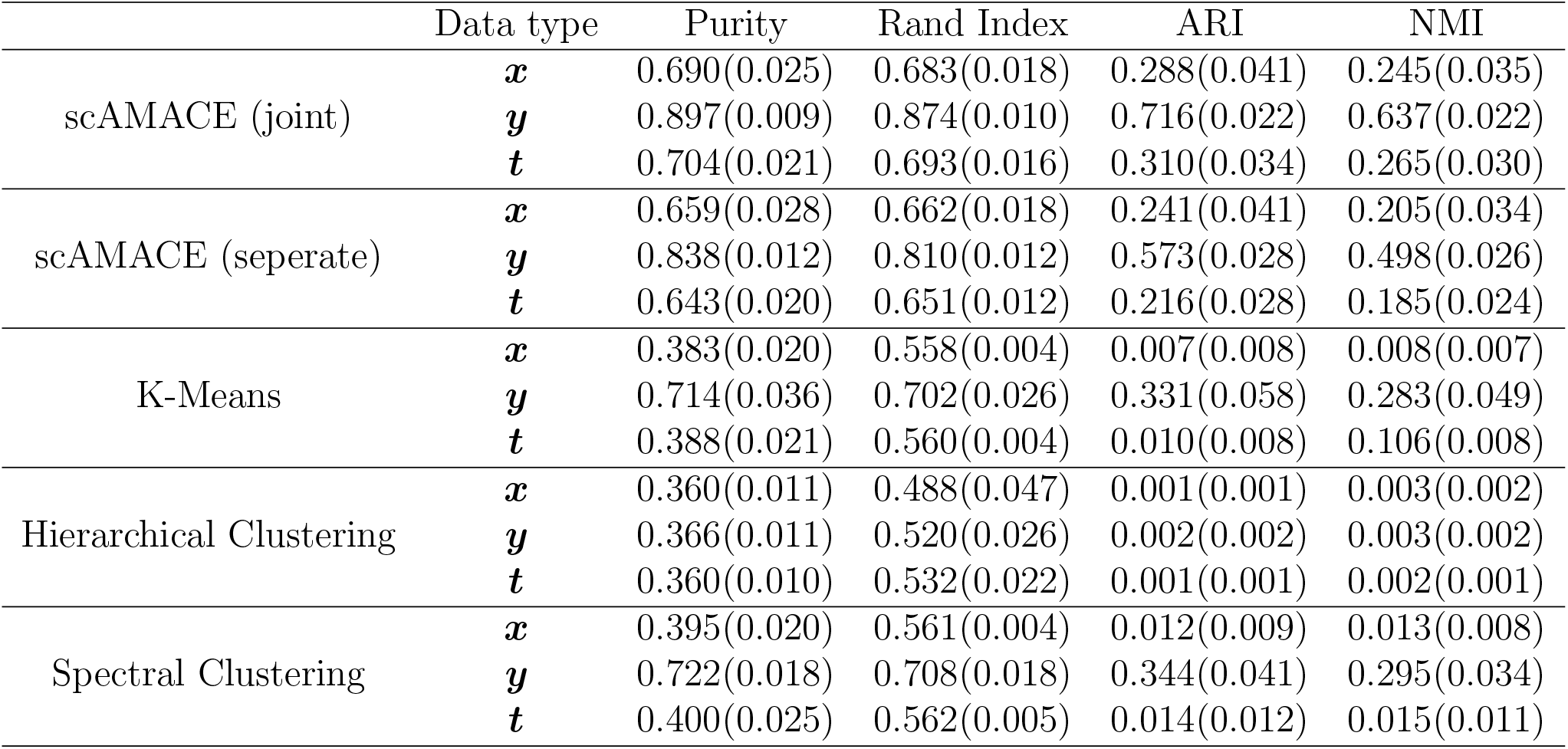
Mean and sd (in parentheses) of purity, rand index, adjusted rand index (ARI) and normalized mutual information (NMI) for 50 independent runs are shown.

|  | Data type | Purity | Rand Index | ARI | NMI |
| --- | --- | --- | --- | --- | --- |
| scAMACE (joint) | $x$ | 0.690(0.025) | 0.683(0.018) | 0.288(0.041) | 0.245(0.035) |
| | $y$ | 0.897(0.009) | 0.874(0.010) | 0.716(0.022) | 0.637(0.022) |
| | $t$ | 0.704(0.021) | 0.693(0.016) | 0.310(0.034) | 0.265(0.030) |
| scAMACE (seperate) | $x$ | 0.659(0.028) | 0.662(0.018) | 0.241(0.041) | 0.205(0.034) |
| | $y$ | 0.838(0.012) | 0.810(0.012) | 0.573(0.028) | 0.498(0.026) |
| | $t$ | 0.643(0.020) | 0.651(0.012) | 0.216(0.028) | 0.185(0.024) |
| K-Means | $x$ | 0.383(0.020) | 0.558(0.004) | 0.007(0.008) | 0.008(0.007) |
| | $y$ | 0.714(0.036) | 0.702(0.026) | 0.331(0.058) | 0.283(0.049) |
| | $t$ | 0.388(0.021) | 0.560(0.004) | 0.010(0.008) | 0.106(0.008) |
| Hierarchical Clustering | $x$ | 0.360(0.011) | 0.488(0.047) | 0.001(0.001) | 0.003(0.002) |
| | $y$ | 0.366(0.011) | 0.520(0.026) | 0.002(0.002) | 0.003(0.002) |
| | $t$ | 0.360(0.010) | 0.532(0.022) | 0.001(0.001) | 0.002(0.001) |
| Spectral Clustering | $x$ | 0.395(0.020) | 0.561(0.004) | 0.012(0.009) | 0.013(0.008) |
| | $y$ | 0.722(0.018) | 0.708(0.018) | 0.344(0.041) | 0.295(0.034) |
| | $t$ | 0.400(0.025) | 0.562(0.005) | 0.014(0.012) | 0.015(0.011) |

**Table 2:** Clustering tables for K562, GM12878 scRNA-Seq, scATAC-Seq and sc-methylation data.

|  |  | scAMACE (joint) |  |  | scAMACE (seperate) |  |  |
| --- | --- | --- | --- | --- | --- | --- | --- |
|  |  | 1 | 2 |  | 1 | 2 |  |
| scATAC-Seq | GM12878 | 368 | 5 |  | 254 | 119 |  |
|  | K562 | 6 | 660 |  | 171 | 495 |  |
| scRNA-Seq | GM12878 | 128 | 0 |  | 128 | 0 |  |
|  | K562 | 0 | 73 |  | 0 | 73 |  |
| sc-methyl | GM12878 | 16 | 3 |  | 7 | 12 |  |
|  | K562 | 0 | 11 |  | 11 | 0 |  |
|  |  | Seurat V3 |  |  | LIGER |  |  |
|  |  | 1 | 2 | 3 | 1 | 2 | 3 |
| scATAC-Seq | GM12878 | 346 | 27 |  |  |  |  |
|  | K562 | 499 | 167 |  |  |  |  |
| scRNA-Seq | GM12878 | 101 | 2 | 25 | 127 | 0 | 1 |
|  | K562 | 0 | 73 | 0 | 10 | 63 | 0 |
| sc-methyl | GM12878 |  |  |  |  |  | 19 |
|  | K562 |  |  |  |  |  | 11 |

**Table 3:** Comparison of the performance of different methods on the K562, GM12878 dataset by adjusted rand index (ARI).

|  | scAMACE (joint) | scAMACE (seperate) | Seurat V3 | LIGER | scMC |
| --- | --- | --- | --- | --- | --- |
| scATAC-Seq | 0.958 | 0.192 | 0.033 |  | 0.000 |
| scRNA-Seq | 1.000 | 1.000 | 0.713 | 0.800 | 0.771 |
| sc-methyl | 0.628 | 0.260 |  | 0.000 | 0.000 |

**Table 4:**
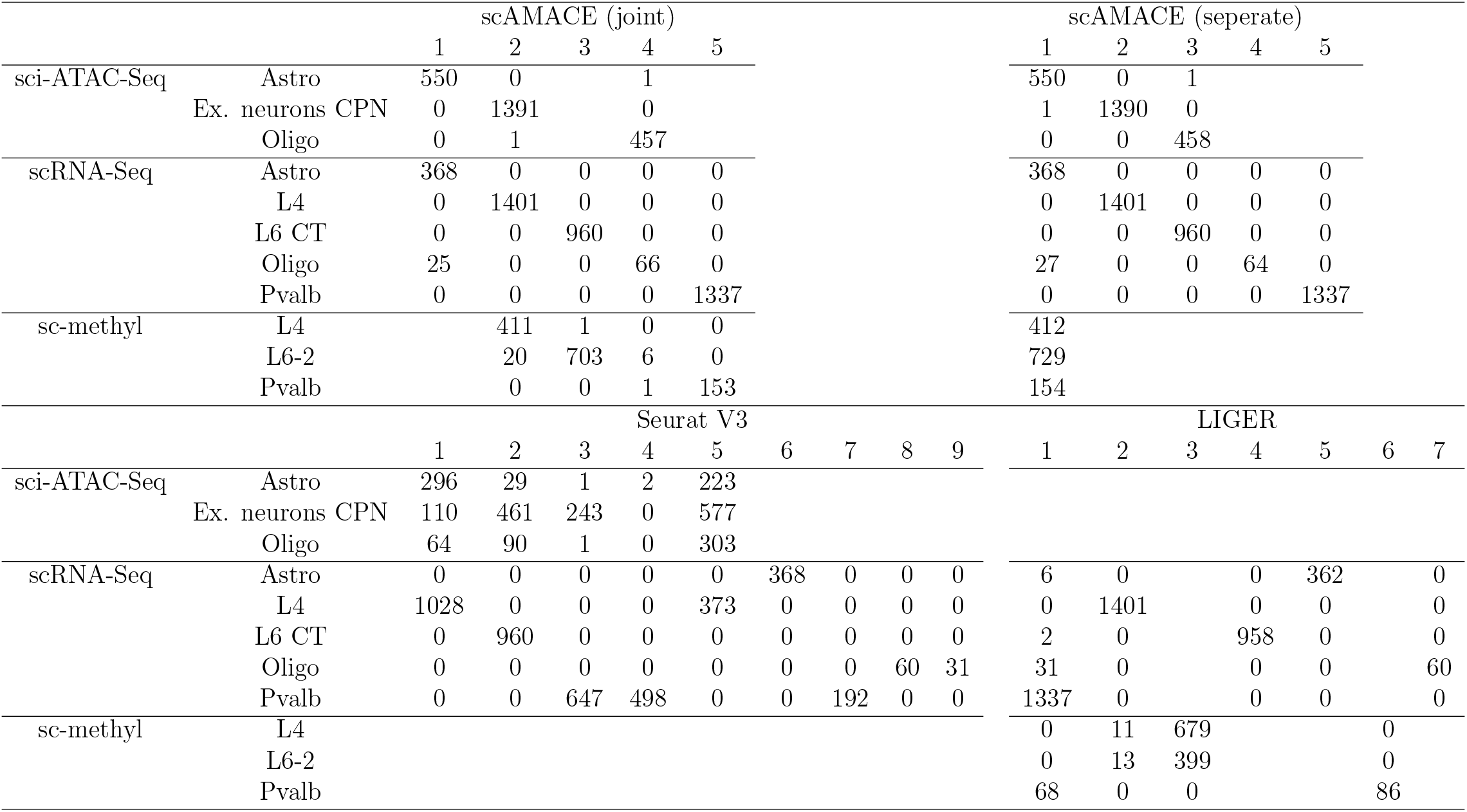
Clustering tables for the mouse neocortex scRNA-Seq, sci-ATAC-Seq, and sc-methylation data.

**Table 5:**
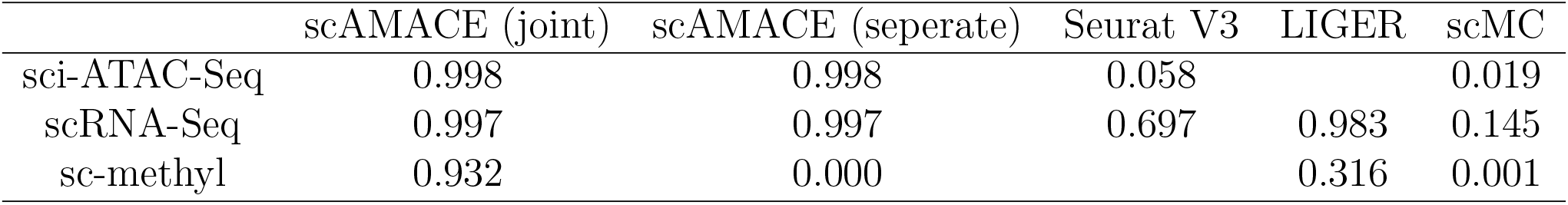
Comparison of the performance of different methods on the mouse neocortex dataset by adjusted rand index (ARI).

In the following two real data applications, we apply methods mentioned in Section 2.7 instead of fitting a beta mixture model to sc-methylation data.

## 5 Application to real data

### 5.1 Application 1: K562 and GM12878 scRNA-Seq, scATAC-Seq and sc-methylation data

We evaluate scAMACE by jointly clustering scRNA-Seq, scATAC-Seq and sc-methylation data generated from two cell types, K562 and GM12878 (Buenrostro *et al*., 2015; Li *et al*., 2017; Pott, 2017). We set *K* = 2, and use the true cell labels as a benchmark to evaluate the performance of the clustering methods. Table S.5 presents the clustering results. scAMACE performs well in seperating the cell types. scRNA-Seq is perfectly seperated, while there are only three cells that are not classified correctly in the sc-methylation dataset and eleven misclassifications in the scATAC-Seq dataset. In addition, the two cell types are correctly matched across the three datasets. Compared with the clustering results given by implementing scAMACE seperately on the three datasets, jointly clustering the three datasets improves the overall clustering performance, especially for scATAC-Seq data, which is likely due to the integration of information across the three datasets.

We compared scAMACE with Seurat V3 (Stuart *et al*., 2019), LIGER (Welch *et al*., 2019) and scMC (Zhang and Nie, 2021), which are methods for integrative analysis of single-cell data. Examples were presented in Seurat V3 (Stuart *et al*., 2019) where scRNA-Seq and scATAC-Seq data were integrated. So we implemented Seurat V3 to integrate these two data types. Seurat V3 did not perform well for scATAC-Seq data (Table S.5). Seurat V3 is not applicable to integrate sc-methylation data with the other two datasets. Examples were presented in LIGER (Welch *et al*., 2019) where scRNA-Seq data and sc-methylation data were integrated. So we implemented LIGER to integrate these two data types. LIGER did not perform well on sc-methylation data (Table S.5). We also implemented LIGER to integrate all three datasets, and LIGER still did not perform well on sc-methylation data (Supplementary Materials Table S.7 and S.8), this may be due to the small sample size in sc-methylation data. scMC (Zhang and Nie, 2021) was developed for the integrative analysis of multiple single-cell datasets with the same data type. Since the features in scATAC-Seq data, scRNA-Seq data and sc-methylation data are linked, scMC can be implemented in principle. scMC did not perform well on scATAC-Seq data and sc-methylation data (Supplementary Materials Table S.7 and S.8). This may be due to the fact that the characteristics of different data types are very different, and ignoring the difference leads to suboptimal performance.

### 5.2 Application 2: Mouse neocortex scRNA-Seq, sci-ATAC-Seq and sc-methylation data

In this example, we evaluate scAMACE for the joint analysis of single-cell datasets where the cell types are different across the datasets.

We collected single-cell datasets generated from mouse neocortex. There are five cell types in scRNA-Seq data (Tasic *et al*., 2018), including astrocytes (Astro), glutamatergic neurons in layer 4 (L4), corticothalamic glutamatergic neurons in layer 6 (L6 CT), oligodendrocytes (Oligo) and Pvalb+ GABAergic neurons (Pvalb). There are three cell types in sci-ATAC-Seq data (Cusanovich *et al*., 2018b), including astrocytes (Astro), excitatory neurons CPN (Ex. neurons CPN), and oligodendrocytes (Oligo). There are three cell types in sc-methylation dataset (Luo *et al*., 2017), including excitatory neurons in layer 4 (L4), excitatory neurons in layer 6 (labeled as L6-2 in (Luo *et al*., 2017)), and Pvalb+ GABAergic neurons (Pvalb). In the three datasets, the optimal numbers of clusters chosen by the Silhoutte method, 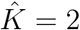, tend to be smaller than the numbers of cell types, which is likely due to the similarity of the neuronal subtypes. We set K=5 when we implement scAMACE, instead of the value given by the Silhoutte method. The true cell labels are used as a benchmark for evaluating the performance of the clustering methods.

The clustering results are presented in Table S.6. Even though *K* is larger than the number of cell types in sci-ATAC-Seq data and sc-methylation data, scAMACE still determines the correct number of cell types in sci-ATAC-Seq data. Although the cells in sc-methylation data fall into four clusters, there are only seven cells in cluster 4. Cell types in all three datasets are well seperated. Astrocytes and oligodendrocytes are matched across scRNA-Seq data and sci-ATAC-Seq data. Excitatory neurons CPN in sci-ATAC-Seq data are matched with glutamatergic neurons in layer 4 in the scRNA-Seq data. We note that most excitatory neurons are glutamatergic neurons. Excitatory neurons in layers 4 and 6, and Pvalb+ GABAergic neurons are matched between scRNA-Seq data and sc-methylation data.

Compared with implementing scAMACE on the three datasets separately, the joint analysis leads to improvement in clustering, especially for sc-methylation dataset. This is likely because the joint model borrows information across the three datasets. Similar to application 1, we implemented Seurat V3 to integrate scRNA-Seq and sci-ATAC-Seq data. Seurat V3 (Stuart *et al*., 2019) does not perform well on sci-ATAC-Seq data (Table S.6). We implemented LIGER (Welch *et al*., 2019) to integrate scRNA-Seq and sc-methylation data. LIGER does not seperate excitatory neurons in layer 4 and layer 6 in sc-methylation data (Table S.6). We also integrated all three datasets by LIGER (Welch *et al*., 2019) and scMC (Zhang and Nie, 2021). LIGER and scMC did not perform well (Supplementary Materials Table S.9 and S.10). Overall, scAMACE performed the best compared with the other methods.

### 5.3 Computational cost

LIGER, Seurat V3 and scMC only provide the versions that are implemented on CPU, while scAMACE can be implemented on both CPU and GPU. We summarized the computational time for scAMACE (CPU version and GPU version in python), LIGER (Welch *et al*., 2019), Seurat V3 (Stuart *et al*., 2019) and scMC (Zhang and Nie, 2021) (Supplementary Materials Tables S.11, S.12 and S.13). We implemented scAMACE, LIGER and scMC to cluster the three types of data simultaneously, and we implemented Seurat V3 to cluster scCAS data and scRNA-Seq data.

On real data application 2 (∼ 8,000 cells), the computational time for scAMACE are 418.858 seconds on one 3.4GHz Intel Xeon Gold CPU and 69.652 seconds on one 3.1GHz Dual Intel Xeon Gold GPU. Compared with LIGER (80.389 seconds on one 3.4GHz Intel Xeon Gold CPU), scMC (372.323 seconds on one 3.4GHz Intel Xeon Gold CPU) and Seurat V3 (116.688 seconds for scRNA-Seq and sci-ATAC-Seq data on one 3.4GHz Intel Xeon Gold CPU), scAMACE has competitive computational speed, especially the GPU version.

Next, we generated a dataset with sample size=30,000 (*n*_*acc*_ = *n*_*rna*_ = *n*_*met*_ = 10, 000) by sampling the cells with replacement from real data application 2. The computational time for scAMACE are 1534.631 seconds on one 3.4GHz Intel Xeon Gold CPU and 250.089 seconds on one 3.1GHz Dual Intel Xeon Gold GPU. Compared with LIGER (555.574 seconds on one 3.4GHz Intel Xeon Gold CPU), scMC (3667.878 seconds on one 3.4GHz Intel Xeon Gold CPU) and Seurat V3 (290.640 seconds for scRNA-Seq and sci-ATAC-Seq data on one 3.4GHz Intel Xeon Gold CPU), scAMACE has competitive computational speed on datasets with larger scale.

## 6 Conclusion

Unsupervised methods including dimension reduction and clustering are essential to the analysis of single-cell genomic data as the cell types are usually unknown. We have developed scAMACE, a model-based approach for integratively clustering single-cell data on chromatin accessibility, gene expression and methylation. scAMACE provides statistical inference of cluster assignments and achieves better cell type seperation combining biological information across different types of genomic features. In the two real data applications, the scRNA-Seq data are generated from the SMART-Seq platform (Li *et al*., 2017; Tasic *et al*., 2018). To implement scAMACE on UMI-based scRNA-Seq data (10x data), we may need to modify the distributions of the mixture components *g*_0_(·) and *g*_1_(·). The cells in our real data examples are differentiated and mature cells. In the future, we will investigate the performance of scAMACE on immature cells undergoing differentiation.

## Acknowledgements

We would like to thank Jinwen Yang, Wenyu Zhang and Pengcheng Zeng for the helpful discussions.

## SUPPLEMENTARY MATERIALS

### S.1 Supplementary Text

#### S.1.1 Joint likelihood

- scCAS data

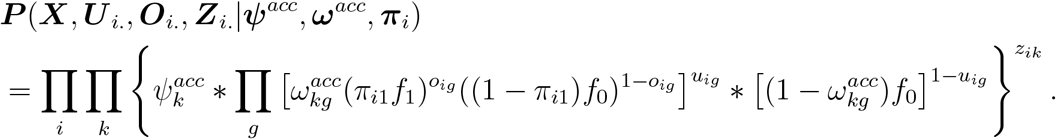
- scRNA-Seq data

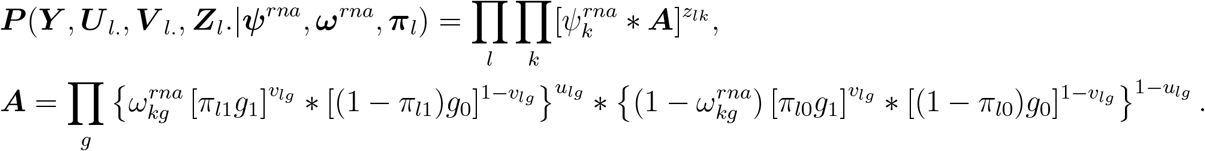
- sc-methylation data

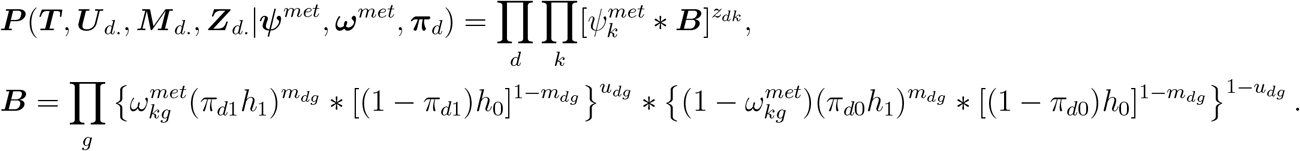

#### S.1.2 Q-function

Let **Γ** denote the missing data, and let **Φ** denote the parameters. the Q-function is ***Q***(**Φ**|**Φ**_*old*_) = 𝔼_*old*_(*ln*(***P*** (**Φ, Γ**|*obs*.))), where the expectation is over **Γ** under distribution ***P*** (**Γ**|**Φ**^*old*^, *obs*.) := ***P*** _*old*_(**Γ**).

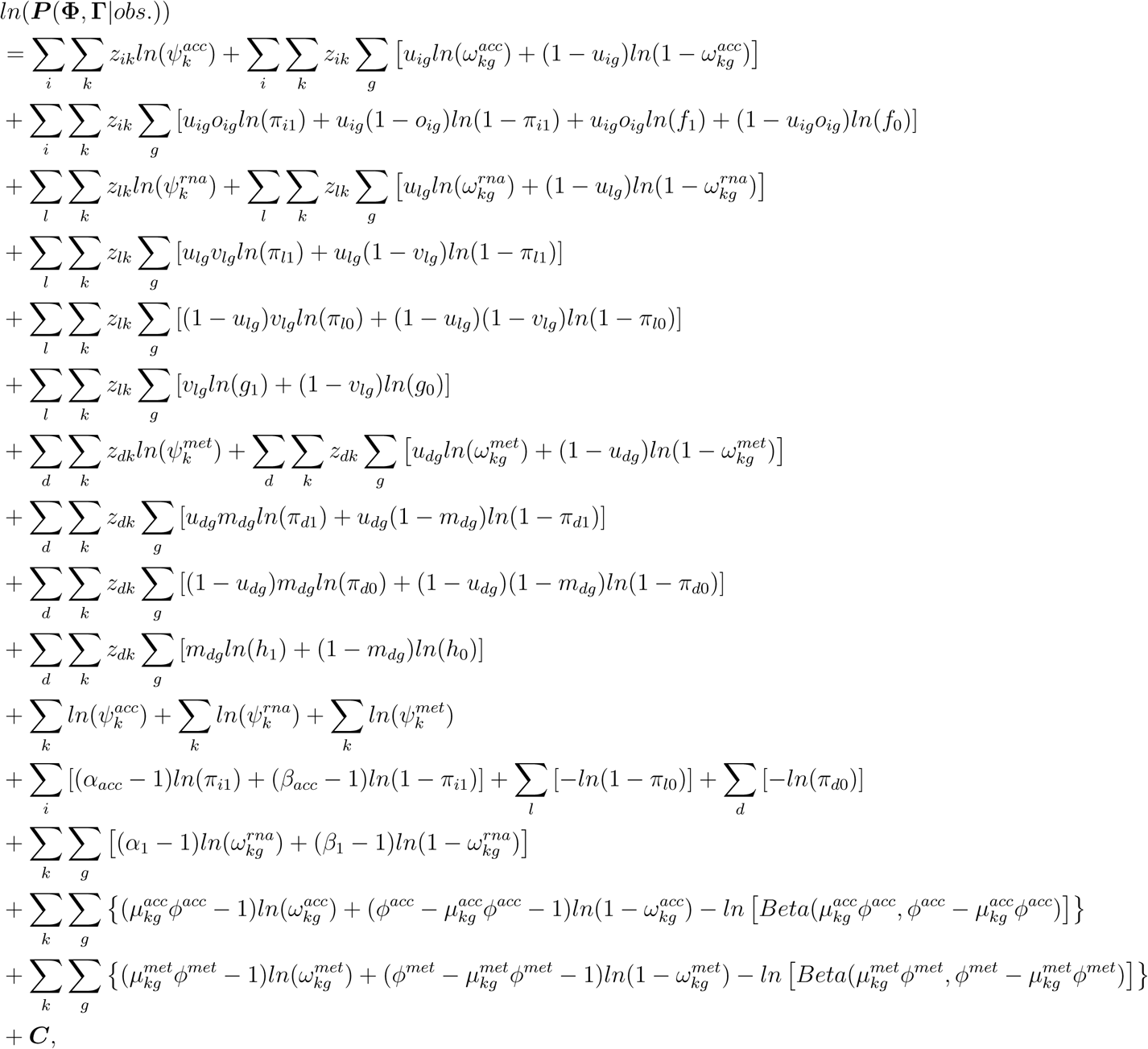

where 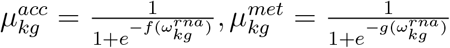, and ***C*** is a constant that does not depend on the parameters.

#### S.1.3 Expectations in E-Step

- scCAS data

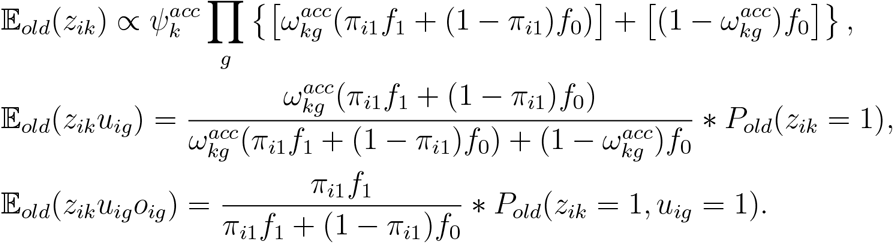
- scRNA-Seq data

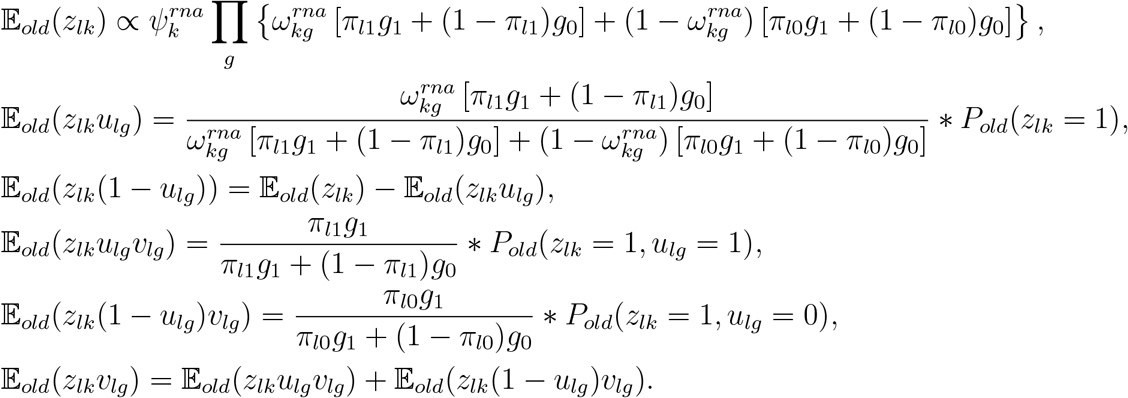
- sc-methylation data

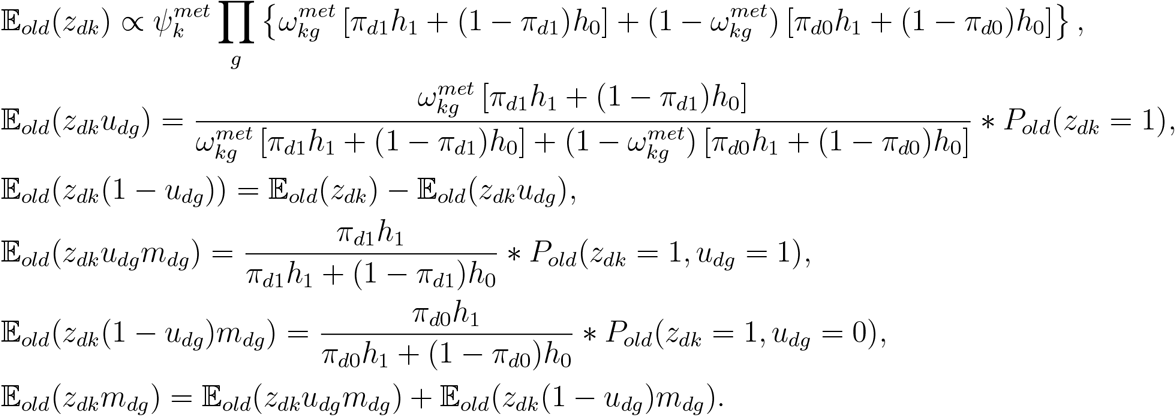

#### S.1.4 Simulation scheme

We generated three different types of simulated data ***x, y*** and ***t*** following the model assumption. In the simulated data, the sample sizes *n*_*x*_ = 900, *n*_*y*_ = 1100, and *n*_*t*_ = 1000. The number of features *p* = 1000. The numbers of clusters *K*_*x*_ = *K*_*y*_ = *K*_*t*_ = 3. 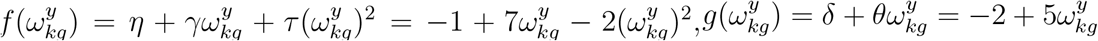, *Φ*^*x*^ = 10 and *Φ*^*t*^ = 10. The followings are the simulation scheme:

##### A. Generate ***ω***^*y*^

For *g* = 1, …, 150:

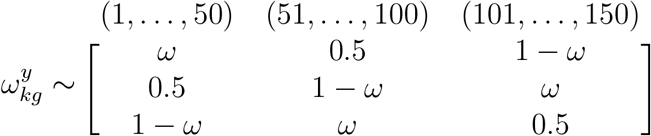

We set *ω* = 0.8.

For *g* = 151, …, 1000:

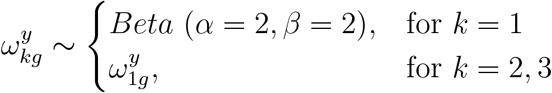

To summarize, we set the first 150 features to be differential and for the remaining 151, …, *p* features, we set 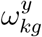 to be the same across different clusters *k*.

##### B. Generate ***ω***^*x*^ and ***ω***^*t*^

For *g* = 1, …, 150:

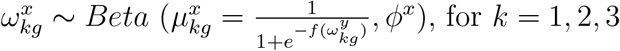

For *g* = 151, …, 1000:

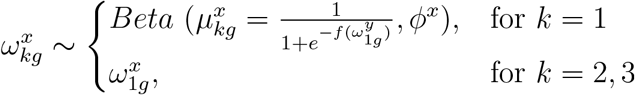

For *g* = 1, …, 150:

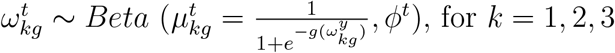

For *g* = 151, …, 1000:

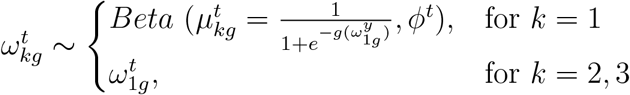

##### C. Generate ***z***^*x*^, ***z***^*y*^ and ***z***^*t*^

The cluster labels are generated with equal probability, 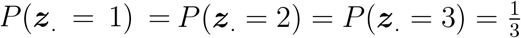.

##### D. Data type 1: ***x***

- Generate ***u***^*x*^. We generate *u*_*ig*_ from *Bernoulli*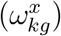 if *z*_*ik*_ = 1.
- Generate ***o***^*x*^. We generate *o*_*ig*_ from *Bernoulli*(*π*_*i*1_) if *u*_*ig*_ = 1, and set *o*_*ig*_ = 0 if *u*_*ig*_ = 0. We set *π*_*i*1_ = 0.2 for *i* = 1, · · ·, *n*_*x*_.
- Generate ***x***. We generate *x*_*ig*_ = 1 if *o*_*ig*_ = 1, and generate *x*_*ig*_ = 0 if *o*_*ig*_ = 0.

##### E. Data type 2: ***y***

- Generate ***u***^*y*^. We generate *u*_*lg*_ from *Bernoulli*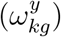 if *z*_*lk*_ = 1.
- Generate ***v***^*y*^. We generate *v*_*lg*_ from *Bernoulli*(*π*_*l*1_) if *u*_*lg*_ = 1, and from *Bernoulli*(*π*_*l*0_) if *u*_*lg*_ = 0. We set *π*_*l*1_ = 0.7, *π*_*l*0_ = 0.3 for *l* = 1, · · ·, *n*_*y*_.
- Generate ***y***. We generate *y*_*lg*_ from *Gamma*(*shape* = 7, *scale* = 0.5) if *v*_*lg*_ = 1, and generate *y*_*lg*_ from *Gamma*(*shape* = 1, *scale* = 1) if *v*_*lg*_ = 0.

##### F. Data type 3: *t*

- Generate ***u***^*t*^. We generate *u*_*dg*_ from *Bernoulli*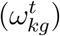 if *z*_*dk*_ = 1.
- Generate ***m***^*t*^. We generate *m*_*dg*_ from *Bernoulli*(*π*_*d*1_) if *u*_*dg*_ = 1, and from *Bernoulli*(*π*_*d*0_) if *u*_*dg*_ = 0. We set *π*_*d*1_ = 0.4, *π*_*d*0_ = 0.7 for *d* = 1, · · ·, *n*_*t*_.
- Generate ***t***. We generate *t*_*dg*_ from *Beta*(*α* = 0.5, *β* = 0.5) if *m*_*dg*_ = 1, and generate *t*_*dg*_ from *Beta*(*α* = 1, *β* = 10) if *m*_*dg*_ = 0.

We set different parameter values for the four additional simulation settings mentioned in Section 4 as following:

1. Simulation setting 1: imbalanced dataset, where the numbers of cells, *n*_*x*_, *n*_*y*_, and *n*_*t*_ are different across modalities (Table S.1). Data is generated as described above, but we set *n*_*x*_ = 1000, *n*_*y*_ = 2000, and *n*_*t*_ = 500.
2. Simulation setting 2: unequal number of clusters, where the numbers of clusters in the three modalities, *K*_*x*_, *K*_*y*_, *K*_*t*_ are different (Table S.2). Data is generated as described above, but we applied following scheme to generate ***ω***^*y*^ and ***ψ***_*x*_, ***ψ***_*y*_, ***ψ***_*t*_ so that *K*_*x*_ = 3, *K*_*y*_ = 7, *K*_*t*_ = 5.

A. Generate ***ω***^*y*^ For *g* = 1, …, 350:

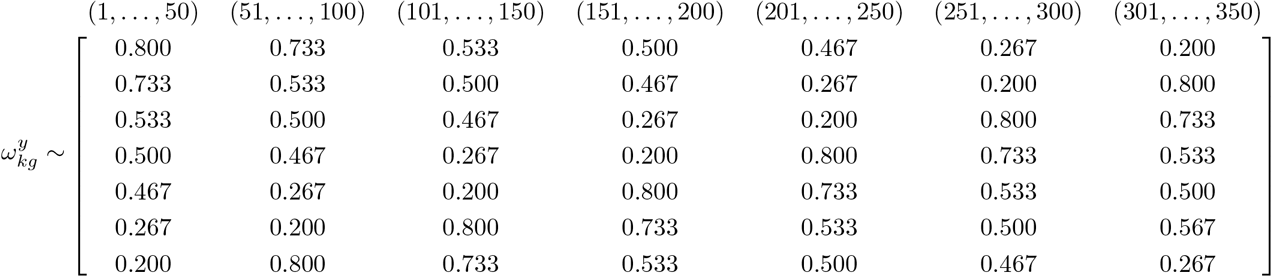 For *g* = 351, …, 1000:

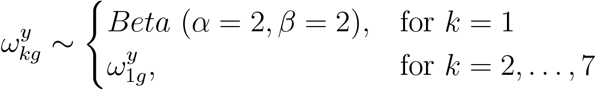
B. Generate ***ω***^*x*^ and ***ω***^*t*^. For *g* = 1, …, 350:

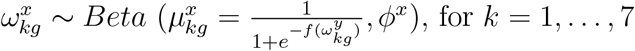 For *g* = 351, …, 1000:

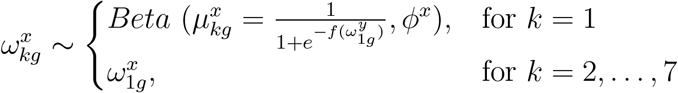 For *g* = 1, …, 350:

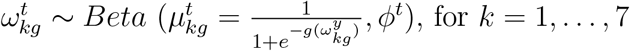 For *g* = 351, …, 1000:

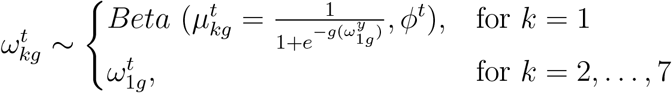
C. Generate ***z***^*x*^, ***z***^*y*^ and ***z***^*t*^. The cluster labels are generated by 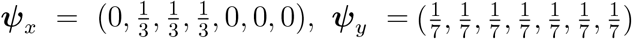, and 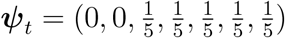 so that *K*_*x*_ = 3, *K*_*y*_ = 7, *K*_*t*_ = 5.

The remaining steps are the same as described above.

(3) Simulation setting 3: imbalanced cluster sizes, where the proportions of different cell types in the three modalities, ***ψ***_*x*_, ***ψ***_*y*_, ***ψ***_*t*_ are different (Table S.3). Data is generated as described above, but we set ***ψ***_*x*_ = (0.3, 0.1, 0.6), ***ψ***_*y*_ = (0.6, 0.3, 0.1), ***ψ***_*t*_ = (0.6, 0.1, 0.3). There are rare cell types (10% of the cells) in the three modalities.
(4) Simulation setting 4: smaller number of features (Table S.4). Data is generated as described above, but we set *p* = 500.

### S.2 Supplementary Figures

#### S.2.1 Histograms for two real data applications

**Figure S.1:**
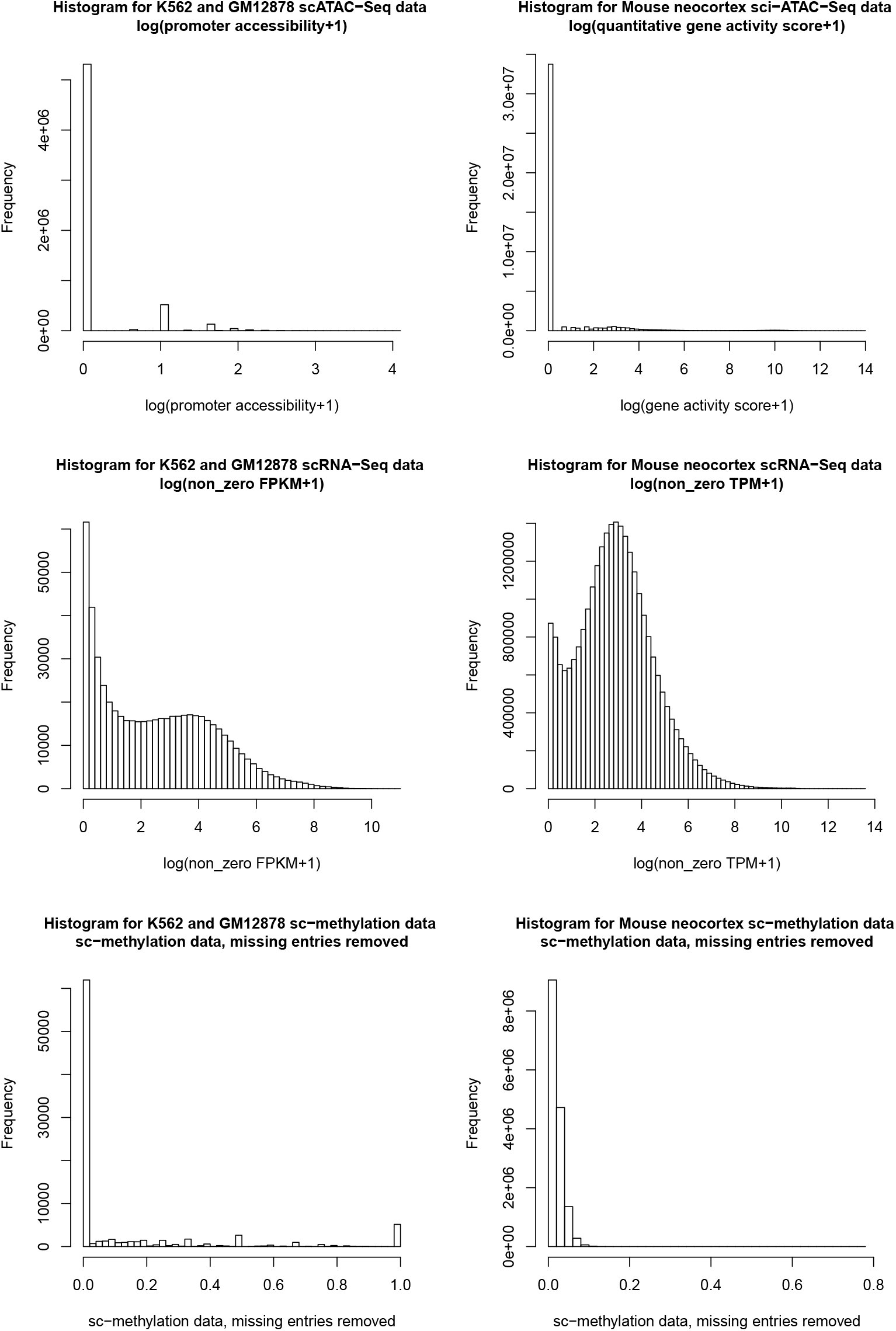
Histograms for Application 1 (Left) and Appliation 2 (Right) scCAS data (Upper), scRNA-Seq data (Middle) and sc-methylation data(Lower).

#### S.2.2 Determination of number of clusters *K* for real data applications

We applied the Silhouette method (Kaufman and Rousseeuw, 1990) mentioned in Section 2.9 on the two real data applications to determine *K* before we apply scAMACE. The result for real application 1 is presented in Figure S.2: 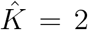 is chosen for the three single-cell datasets, where the true number of cell types is 2. The results for real data application 2 are presented in Figure S.3. There are five cell types in scRNA-Seq data (Tasic *et al*., 2018), including astrocytes, oligodendrocytes, and three subtypes of neurons. There are three cell types in sci-ATAC-Seq data (Cusanovich *et al*., 2018b), including astrocytes, oligodendrocytes, and excitatory neurons CPN. And there are three cell types in sc-methylation dataset (Luo *et al*., 2017), including three subtypes of neurons. In the three datasets, the optimal numbers of clusters chosen by the Silhoutte method 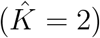 tend to be smaller than the numbers of cell types, which is likely due to the similarity of the neuronal subtypes. We chose *K* = 5 when we implement scAMACE, instead of the suggested 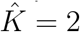 by the Silhoutte method.

**Figure S.2:**
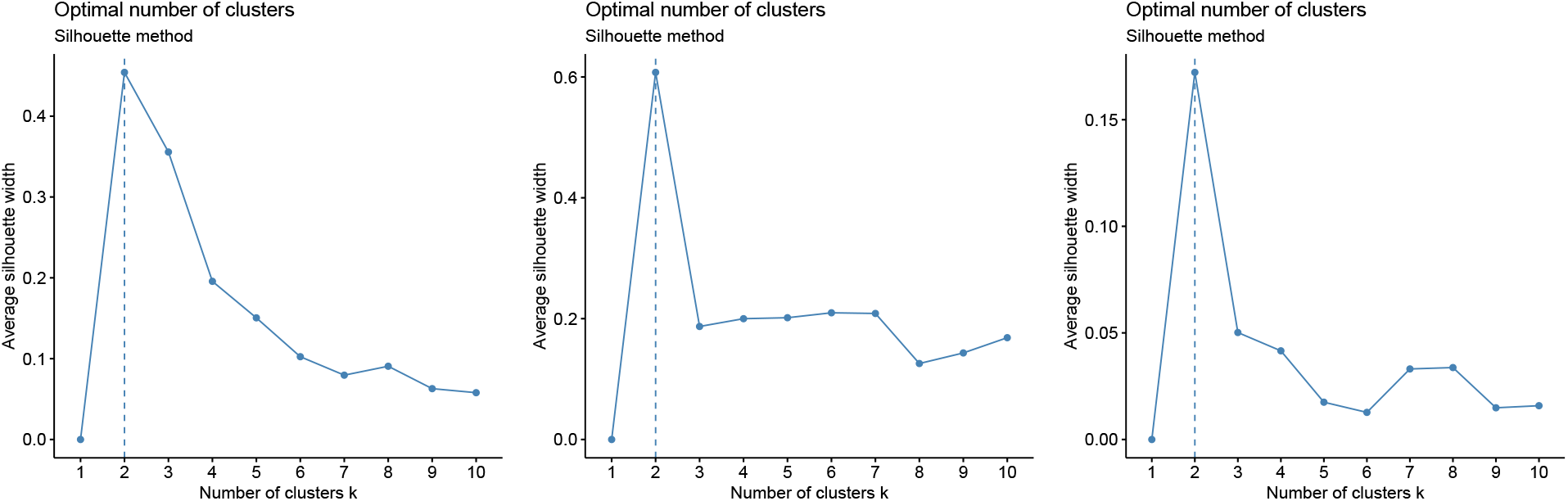
Average Silhouette width v.s. *K* for Application 1, K562 and GM12878 cells: scATAC-Seq (Left), scRNA-Seq (Middle), and sc-methylation data (Right).

**Figure S.3:**
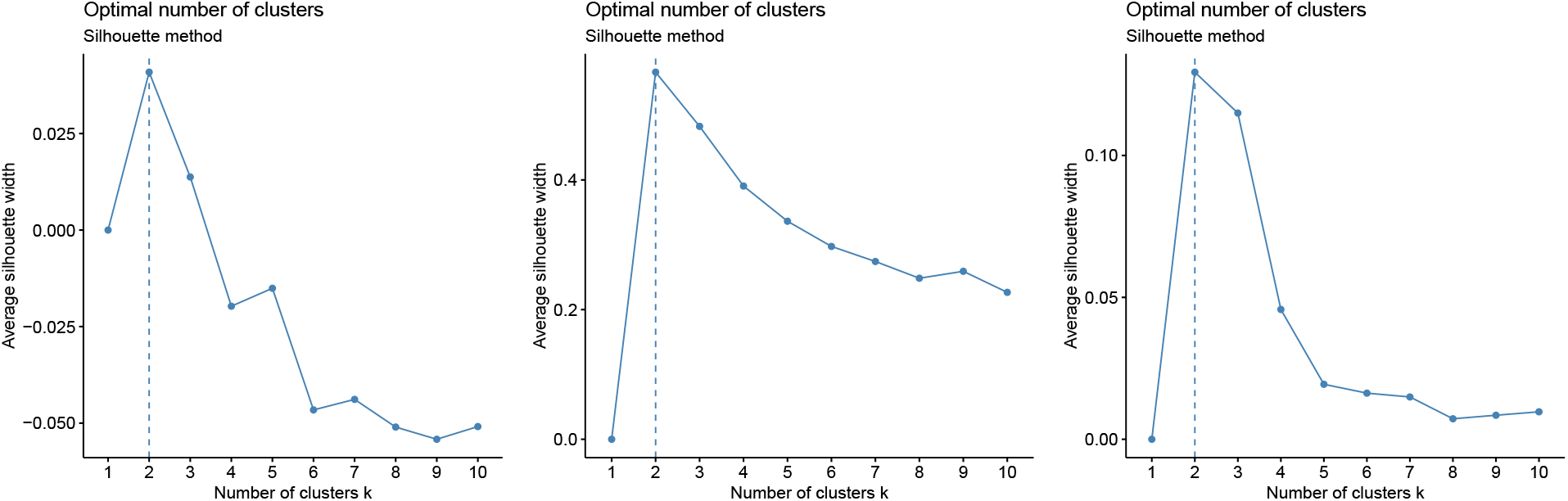
Average Silhouette width v.s. *K* for Application 2, mouse neocortex data: sci-ATAC-Seq (Left), scRNA-Seq (Middle), and sc-methylation data (Right).

#### S.2.3 Empirical distribution of *ω*

To assess whether the linear and quadratic models work well, we performed the following steps. We first obtain 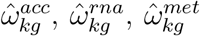 as described in Section 2.6 by setting *K* = 1 and fitting the model on the three modalities separately. We plotted the distributions of 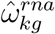 v.s. 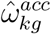 and 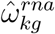 v.s. 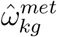 for the two real data applications in the left panels in Figures S.4 and S.5, respectively. For better visualization on how 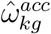 and 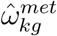 change with 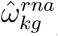, we plotted the boxplots of 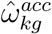 and 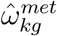 in different ranges of 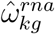. We estimated {*η, γ, τ, δ, θ, ϕ*^*acc*^, *ϕ*^*met*^} by beta regression (Silvia and Francisco, 2004) using 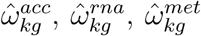. Using 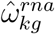 and 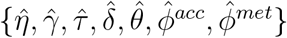, we then generated 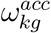 and 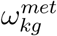 by random sampling following the quadratic and linear models. The distributions of the generated 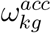 and 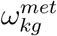 are plotted in the right panels in Figures S.4 and S.5, respectively. In comparison of the generated values (left panels in Figures S.4 and S.5) with the estimated 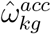 and 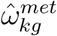 (right panels in Figures S.4 and S.5), we can see that the linear and quadratic models capture the trends on how 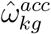 and 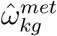 changes with 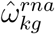.

**Figure S.4:**
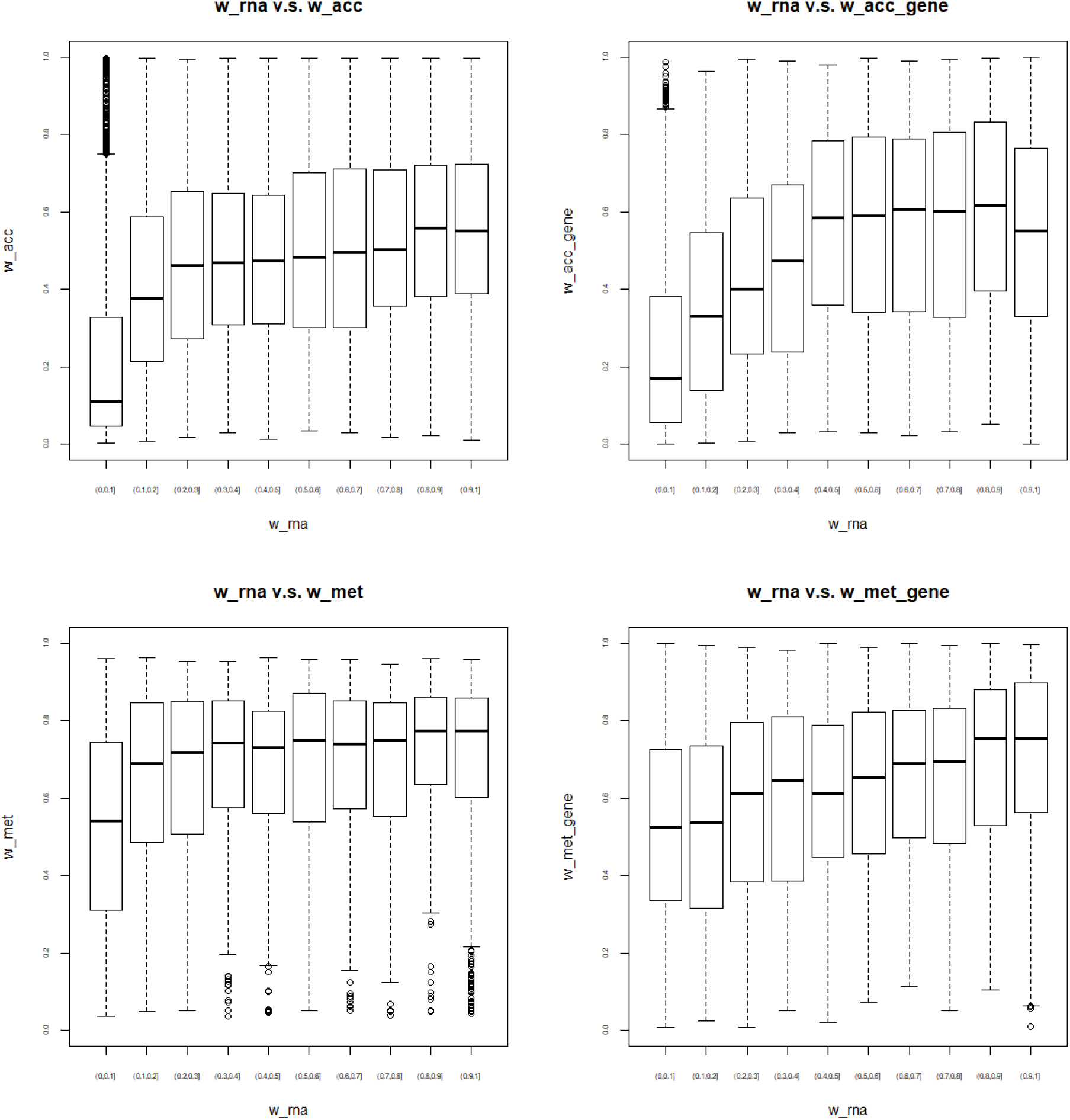
Application 1: Distribution of 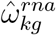 v.s. 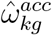 where 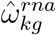 and 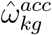 are obtained by setting *K* = 1 and fitting the model on scRNA-Seq data and scCAS data seperately (Top left); Distribution of 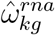 v.s. generated 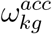 by random sampling from the quadratic model with 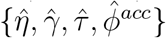 and 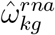 (Top right). Distribution of 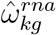 v.s. 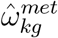 where 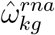 and 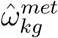 are obtained by setting *K* = 1 and fitting the model on scRNA-Seq data and sc-methylation data seperately (Bottom left); Distribution of 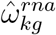 v.s. generated 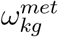 by random sampling from the linear model with 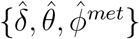 and 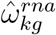 (Bottom right). The estimated values 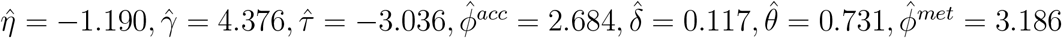.

**Figure S.5:**
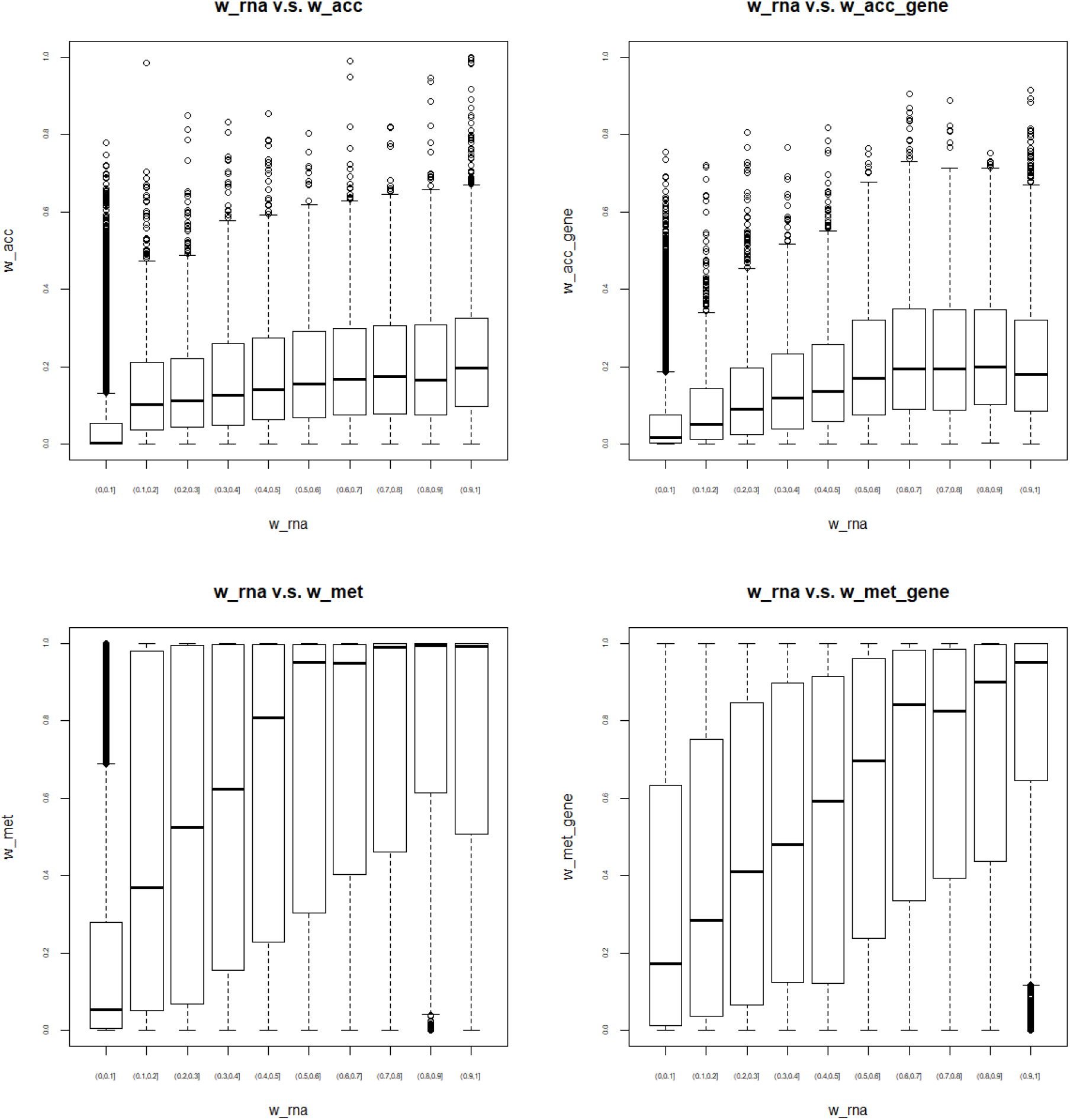
Application 2: Distribution of 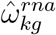 v.s. 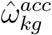 where 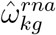 and 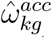 are obtained by setting *K* = 1 and fitting the model on scRNA-Seq data and scCAS data seperately (Top left); Distribution of 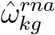 v.s. generated 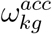 by random sampling from the quadratic model with 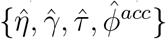 and 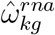 (Top right). Distribution of 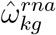 v.s. 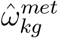 where 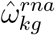 and 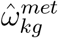 are obtained by setting *K* = 1 and fitting the model on scRNA-Seq data and sc-methylation data seperately (Bottom left); Distribution of 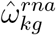 v.s. generated 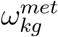 by random sampling from the linear model with 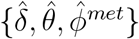 and 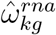 (Bottom right). The estimated values 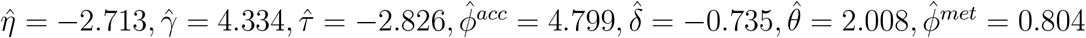.

### S.3 Supplementary Tables

**Table S.1:** Simulation setting 1: imbalanced datasets across the three modalities. Data is generated as described in Section 4, but we set the numbers of cells in the three modalities, *n*_*x*_, *n*_*y*_, and *n*_*t*_ to be different. Mean and sd (in parentheses) of purity, rand index, adjusted rand index (ARI) and normalized mutual information (NMI) for 50 independent runs are shown.

|  |  | Data type | Purity | Rand Index | ARI | NMI |
| --- | --- | --- | --- | --- | --- | --- |
| $n_x = 1000$<br>$n_y = 2000$<br>$n_t = 500$ | scAMACE (joint) | $\mathbf{x}$ | 0.697(0.022) | 0.688(0.017) | 0.299(0.037) | 0.254(0.031) |
| | | $\mathbf{y}$ | 0.911(0.006) | 0.890(0.007) | 0.752(0.016) | 0.673(0.017) |
| | | $\mathbf{t}$ | 0.705(0.025) | 0.693(0.019) | 0.311(0.043) | 0.268(0.035) |
| | scAMACE (seperate) | $\mathbf{x}$ | 0.658(0.023) | 0.661(0.015) | 0.239(0.034) | 0.203(0.029) |
| | | $\mathbf{y}$ | 0.872(0.008) | 0.846(0.008) | 0.652(0.018) | 0.572(0.018) |
| | | $\mathbf{t}$ | 0.650(0.031) | 0.655(0.019) | 0.227(0.043) | 0.196(0.036) |
| | K-Means | $\mathbf{x}$ | 0.382(0.019) | 0.558(0.003) | 0.007(0.007) | 0.008(0.006) |
| | | $\mathbf{y}$ | 0.811(0.011) | 0.784(0.010) | 0.514(0.023) | 0.442(0.021) |
| | | $\mathbf{t}$ | 0.394(0.024) | 0.559(0.005) | 0.009(0.009) | 0.011(0.009) |
| | Hierarchical Clustering | $\mathbf{x}$ | 0.358(0.008) | 0.493(0.045) | 0.001(0.001) | 0.002(0.002) |
| | | $\mathbf{y}$ | 0.356(0.007) | 0.526(0.023) | 0.001(0.001) | 0.002(0.001) |
| | | $\mathbf{t}$ | 0.372(0.014) | 0.524(0.033) | 0.002(0.002) | 0.005(0.004) |
| | Spectral Clustering | $\mathbf{x}$ | 0.390(0.022) | 0.560(0.004) | 0.011(0.009) | 0.012(0.009) |
| | | $\mathbf{y}$ | 0.806(0.010) | 0.779(0.010) | 0.503(0.021) | 0.431(0.019) |
| | | $\mathbf{t}$ | 0.399(0.025) | 0.560(0.005) | 0.010(0.010) | 0.013(0.009) |

**Table S.2:** Simulation setting 2: unequal number of clusters across the three modalities. Data is generated as described in Section 4, but we set the numbers of clusters in the three modalities, *K*_*x*_, *K*_*y*_, *K*_*t*_ to be different. Mean and sd (in parentheses) of purity, rand index, adjusted rand index (ARI) and normalized mutual information (NMI) for 50 independent runs are shown.

|  |  | Data type | Purity | Rand Index | ARI | NMI |
| --- | --- | --- | --- | --- | --- | --- |
| $K_x = 3$<br>$K_y = 7$<br>$K_t = 5$ | scAMACE (joint) | $\mathbf{x}$ | 0.727(0.023) | 0.714(0.017) | 0.353(0.039) | 0.291(0.032) |
| | | $\mathbf{y}$ | 0.775(0.015) | 0.889(0.006) | 0.549(0.024) | 0.559(0.022) |
| | | $\mathbf{t}$ | 0.628(0.019) | 0.773(0.008) | 0.325(0.076) | 0.291(0.020) |
| | scAMACE (seperate) | $\mathbf{x}$ | 0.717(0.020) | 0.705(0.014) | 0.322(0.033) | 0.231(0.024) |
| | | $\mathbf{y}$ | 0.710(0.018) | 0.864(0.007) | 0.446(0.027) | 0.466(0.026) |
| | | $\mathbf{t}$ | 0.591(0.017) | 0.762(0.006) | 0.241(0.020) | 0.236(0.018) |
| | K-Means | $\mathbf{x}$ | 0.416(0.027) | 0.566(0.008) | 0.285(0.016) | 0.024(0.016) |
| | | $\mathbf{y}$ | 0.282(0.024) | 0.767(0.004) | 0.050(0.016) | 0.079(0.020) |
| | | $\mathbf{t}$ | 0.276(0.016) | 0.684(0.002) | 0.190(0.008) | 0.023(0.008) |
| | Hierarchical Clustering | $\mathbf{x}$ | 0.363(0.012) | 0.485(0.050) | 0.372(0.062) | 0.003(0.002) |
| | | $\mathbf{y}$ | 0.188(0.006) | 0.704(0.038) | 0.002(0.001) | 0.012(0.003) |
| | | $\mathbf{t}$ | 0.240(0.009) | 0.637(0.029) | 0.215(0.024) | 0.007(0.002) |
| | Spectral Clustering | $\mathbf{x}$ | 0.434(0.031) | 0.573(0.010) | 0.292(0.017) | 0.039(0.022) |
| | | $\mathbf{y}$ | 0.323(0.022) | 0.774(0.004) | 0.079(0.014) | 0.121(0.018) |
| | | $\mathbf{t}$ | 0.295(0.019) | 0.688(0.003) | 0.197(0.009) | 0.036(0.010) |

**Table S.3:**
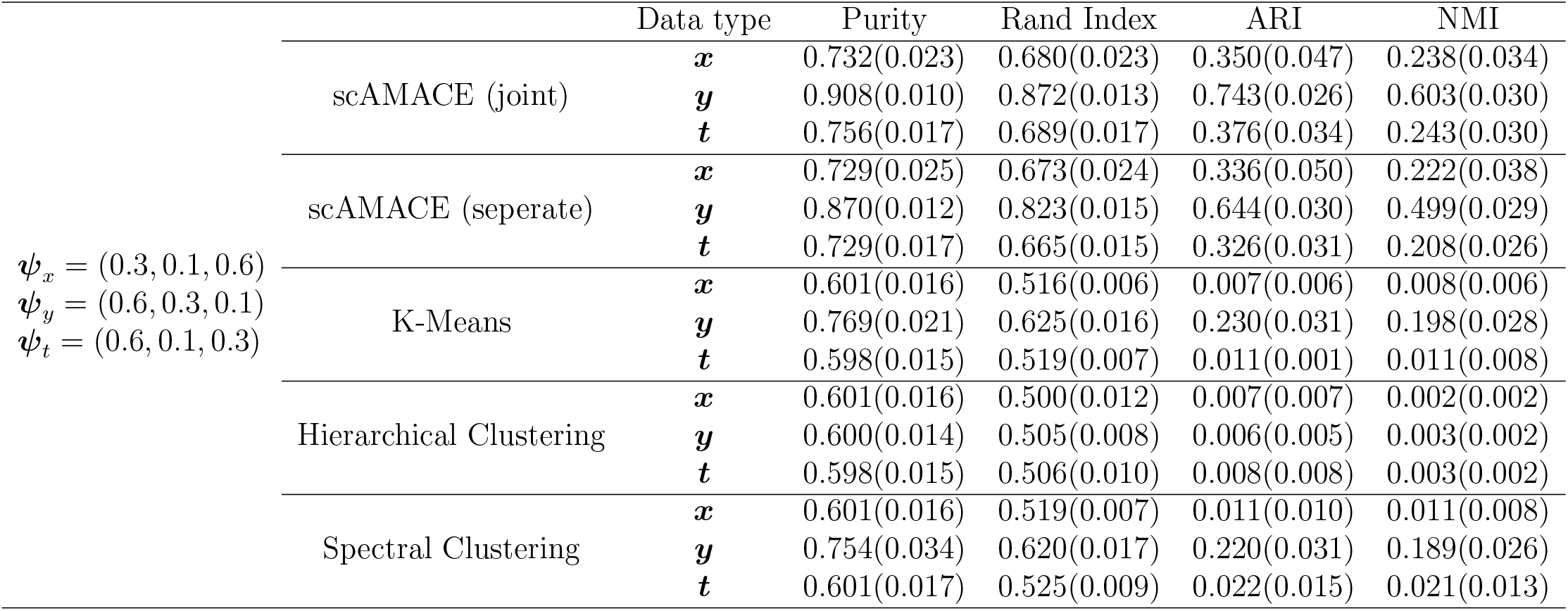
Simulation setting 3: imbalanced cluster sizes across the three modalities. Data is generated as described in Section 4, but we set the proportions different cell types across the three modalities, ***ψ***_*x*_, ***ψ***_*y*_, and ***ψ***_*t*_ to be different. Mean and sd (in parentheses) of purity, rand index, adjusted rand index (ARI) and normalized mutual information (NMI) for 50 independent runs are shown.

**Table S.4:** Simulation setting 4: smaller number of features. Data is generated as described in Section 4, but we set the number of features *p* to be smaller. Mean and sd (in parentheses) of purity, rand index, adjusted rand index (ARI) and normalized mutual information (NMI) for 50 independent runs are shown.

|  |  | Data type | Purity | Rand Index | ARI | NMI |
| --- | --- | --- | --- | --- | --- | --- |
| $p = 500$ | scAMACE (joint) | $\mathbf{x}$ | 0.557(0.029) | 0.605(0.013) | 0.115(0.029) | 0.101(0.025) |
| | | $\mathbf{y}$ | 0.770(0.012) | 0.746(0.010) | 0.429(0.023) | 0.368(0.020) |
| | | $\mathbf{t}$ | 0.564(0.024) | 0.603(0.014) | 0.120(0.025) | 0.107(0.020) |
| | scAMACE (seperate) | $\mathbf{x}$ | 0.507(0.027) | 0.585(0.010) | 0.070(0.022) | 0.063(0.0198) |
| | | $\mathbf{y}$ | 0.658(0.021) | 0.660(0.014) | 0.238(0.030) | 0.205(0.025) |
| | | $\mathbf{t}$ | 0.511(0.023) | 0.581(0.011) | 0.072(0.019) | 0.065(0.016) |
| | K-Means | $\mathbf{x}$ | 0.373(0.013) | 0.557(0.002) | 0.003(0.004) | 0.005(0.004) |
| | | $\mathbf{y}$ | 0.461(0.045) | 0.577(0.013) | 0.049(0.028) | 0.045(0.024) |
| | | $\mathbf{t}$ | 0.377(0.017) | 0.557(0.002) | 0.005(0.005) | 0.006(0.004) |
| | Hierarchical Clustering | $\mathbf{x}$ | 0.359(0.010) | 0.472(0.045) | 0.001(0.001) | 0.003(0.002) |
| | | $\mathbf{y}$ | 0.359(0.009) | 0.501(0.038) | 0.001(0.001) | 0.002(0.002) |
| | | $\mathbf{t}$ | 0.360(0.010) | 0.527(0.023) | 0.001(0.001) | 0.002(0.001) |
| | Spectral Clustering | $\mathbf{x}$ | 0.374(0.015) | 0.557(0.002) | 0.004(0.004) | 0.005(0.004) |
| | | $\mathbf{y}$ | 0.501(0.052) | 0.591(0.015) | 0.079(0.034) | 0.072(0.028) |
| | | $\mathbf{t}$ | 0.379(0.002) | 0.558(0.002) | 0.004(0.004) | 0.006(0.004) |

**Table S.5:** Clustering tables for K562, GM12878 scRNA-Seq, scATAC-Seq and sc-methylation data before and after the transformation on sc-methylation data.

|  |  | scAMACE (joint) |  |  |  |  |  | scAMACE (seperate) |  |  |  |  |  |
| --- | --- | --- | --- | --- | --- | --- | --- | --- | --- | --- | --- | --- | --- |
|  |  | Before transformation |  |  | After transformation |  |  | Before transformation |  |  | After transformation |  |  |
|  |  | 1 | 2 | ARI | 1 | 2 | ARI | 1 | 2 | ARI | 1 | 2 | ARI |
| scATAC-Seq | GM12878 | 368 | 5 | 0.958 | 368 | 5 | 0.958 | 254 | 119 | 0.192 | 254 | 119 | 0.192 |
|  | K562 | 6 | 660 |  | 6 | 660 |  | 171 | 495 |  | 171 | 495 |  |
| scRNA-Seq | GM12878 | 128 | 0 | 1.000 | 128 | 0 | 1.000 | 128 | 0 | 1.000 | 128 | 0 | 1.000 |
|  | K562 | 0 | 73 |  | 0 | 73 |  | 0 | 73 |  | 0 | 73 |  |
| sc-methyl | GM12878 |  | 19 | 0.000 | 16 | 3 | 0.628 |  | 19 | 0.000 | 7 | 12 | 0.260 |
|  | K562 |  | 11 |  | 0 | 11 |  |  | 11 |  | 11 | 0 |  |

**Table S.6:**
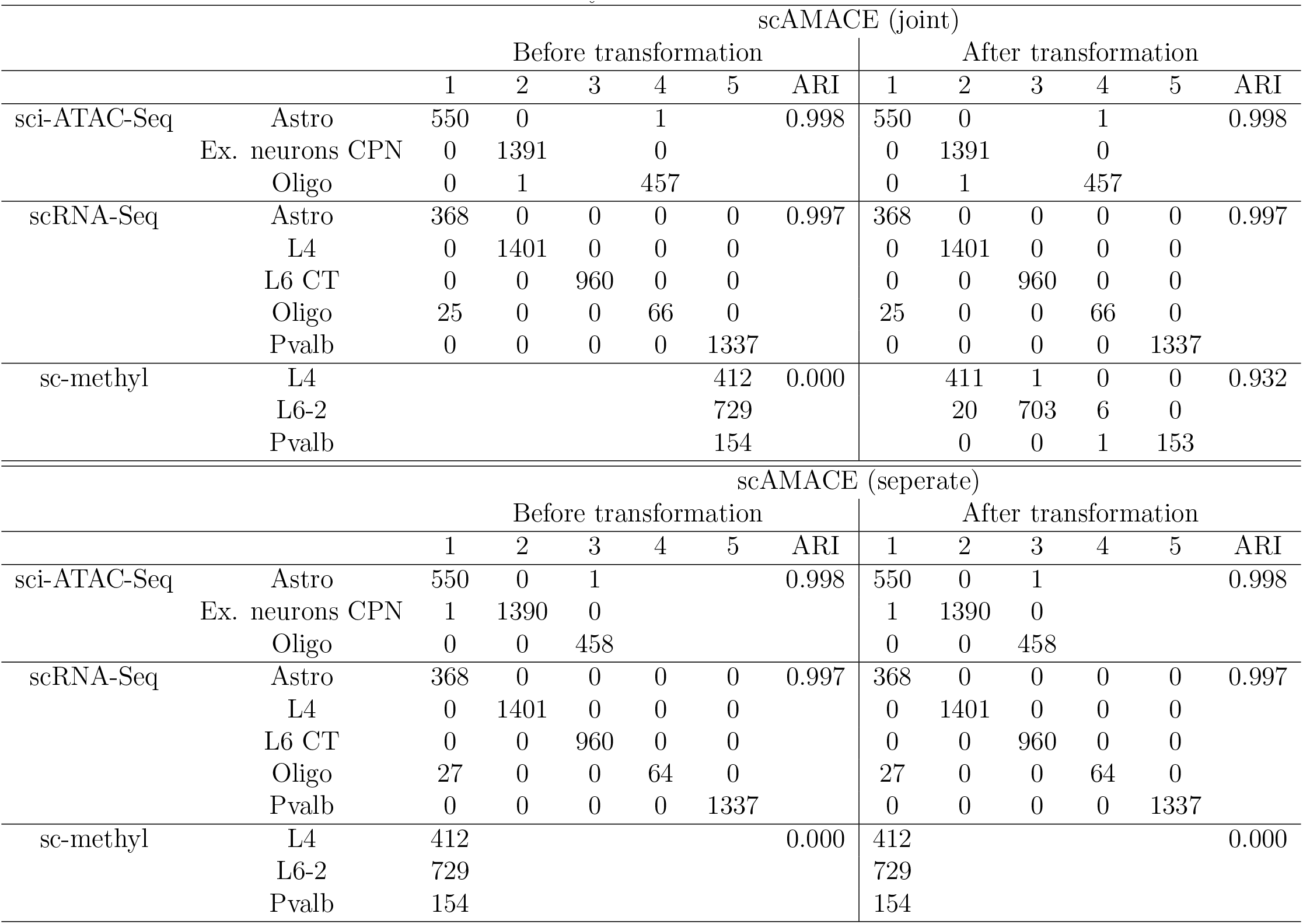
Clustering tables for the mouse neocortex scRNA-Seq, sci-ATAC-Seq, and sc-methylation data before and after the transformation on sc-methylation data.

**Table S.7:**
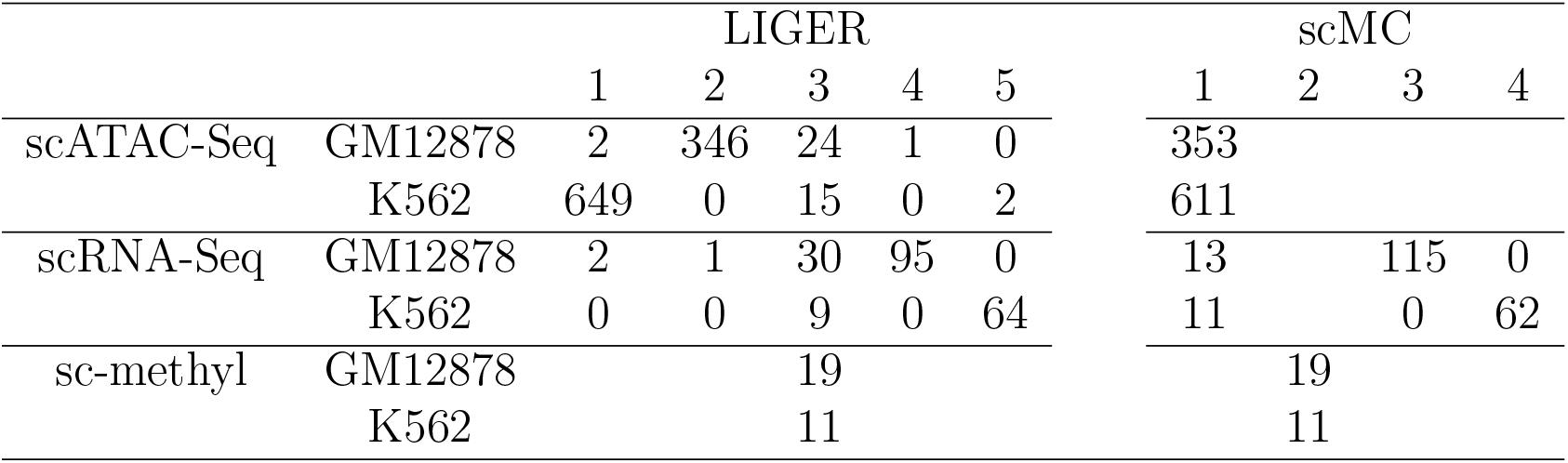
Supplementary clustering tables for K562, GM12878 scRNA-Seq, scATAC-Seq and sc-methylation data.

**Table S.8:** Supplementary comparison of the performance of different methods on the K562, GM12878 dataset by adjusted rand index (ARI).

|  | LIGER | scMC |
| --- | --- | --- |
| scATAC-Seq | 0.918 | 0.000 |
| scRNA-Seq | 0.603 | 0.771 |
| sc-methyl | 0.000 | 0.000 |

**Table S.9:** Supplementary clustering tables for the mouse neocortex scRNA-Seq, sci-ATAC-Seq, and sc-methylation data.

|  |  | LIGER |  |  |  |  |  |  |  |  |  |  |  |
| --- | --- | --- | --- | --- | --- | --- | --- | --- | --- | --- | --- | --- | --- |
|  |  | 1 | 2 | 3 | 4 | 5 | 6 | 7 | 8 | 9 | 10 | 11 | 12 |
| sci-ATAC-Seq | Astro | 145 | 184 | 31 |  | 58 | 12 | 34 | 16 | 10 | 37 | 6 | 0 |
|  | Ex. neurons CPN | 582 | 112 | 194 |  | 135 | 51 | 58 | 87 | 36 | 49 | 3 | 7 |
|  | Oligo | 156 | 62 | 51 |  | 64 | 45 | 25 | 14 | 3 | 13 | 17 | 0 |
| scRNA-Seq | Astro | 49 | 250 | 5 |  | 27 | 5 | 5 | 0 | 8 | 16 | 3 | 0 |
|  | L4 | 1153 | 82 | 66 |  | 1 | 13 | 16 | 13 | 14 | 5 | 38 | 0 |
|  | L6 CT | 791 | 17 | 52 |  | 3 | 40 | 5 | 6 | 27 | 0 | 12 | 7 |
|  | Oligo | 41 | 25 | 3 |  | 3 | 0 | 0 | 2 | 1 | 7 | 9 | 0 |
|  | Pvalb | 1078 | 24 | 78 |  | 3 | 8 | 6 | 9 | 36 | 1 | 22 | 72 |
| sc-methyl | L4 | 137 | 328 | 23 | 174 | 3 | 3 | 6 | 1 | 3 | 6 | 3 | 3 |
|  | L6-2 | 68 | 223 | 18 | 95 | 1 | 1 | 1 | 0 | 3 | 2 | 0 | 0 |
|  | Pvalb | 79 | 27 | 2 | 41 | 2 | 0 | 0 | 0 | 0 | 0 | 1 | 2 |
|  |  | scMC |  |  |  |  |  |  |  |  |  |  |  |
|  |  | 1 | 2 | 3 | 4 | 5 | 6 | 7 |  |  |  |  |  |
| sci-ATAC-Seq | Astro | 8 | 1 | 7 | 326 | 52 | 23 | 133 |  |  |  |  |  |
|  | Ex. neurons CPN | 103 | 18 | 27 | 836 | 164 | 29 | 212 |  |  |  |  |  |
|  | Oligo | 13 | 2 | 10 | 226 | 86 | 16 | 105 |  |  |  |  |  |
| scRNA-Seq | Astro | 0 |  |  | 362 | 4 | 1 | 1 |  |  |  |  |  |
|  | L4 | 1 |  |  | 1 | 1362 | 0 | 37 |  |  |  |  |  |
|  | L6 CT | 0 |  |  | 0 | 959 | 0 | 1 |  |  |  |  |  |
|  | Oligo | 0 |  |  | 14 | 58 | 1 | 18 |  |  |  |  |  |
|  | Pvalb | 1 |  |  | 0 | 1331 | 0 | 5 |  |  |  |  |  |
| sc-methyl | L4 |  |  |  |  | 1 | 689 |  |  |  |  |  |  |
|  | L6-2 |  |  |  |  | 0 | 412 |  |  |  |  |  |  |
|  | Pvalb |  |  |  |  | 0 | 154 |  |  |  |  |  |  |

**Table S.10:** Supplementary comparison of the performance of different methods on the mouse neocorex dataset by adjusted rand index (ARI).

|  | LIGER | scMC |
| --- | --- | --- |
| sci-ATAC-Seq | 0.052 | 0.019 |
| scRNA-Seq | 0.099 | 0.145 |
| sc-methyl | 0.031 | 0.001 |

**Table S.11:**
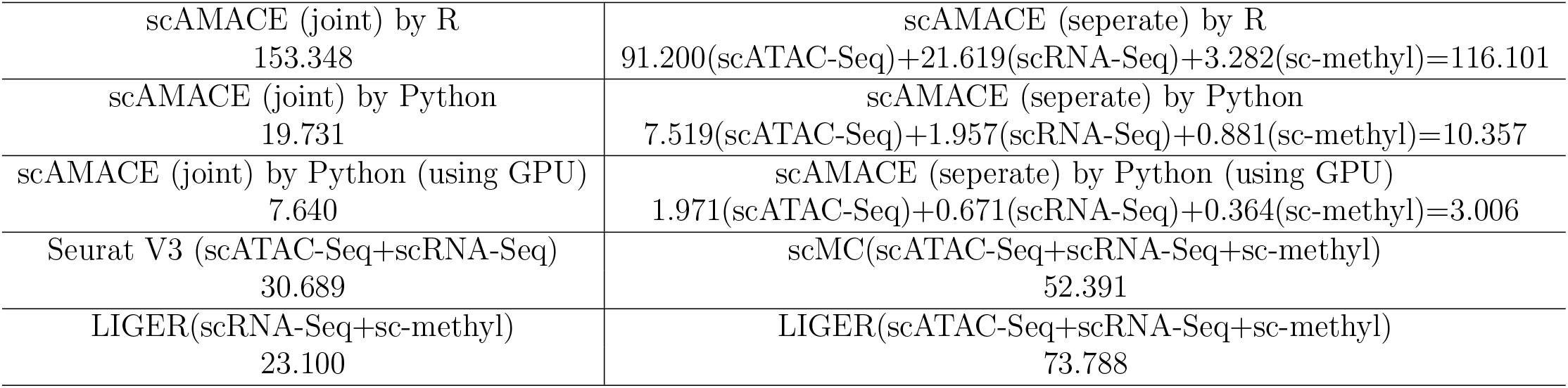
Summary of the computation time by scAMACE and other clustering methods for Application 1. The unit of measurement is second. We implemented scAMACE (jointly on the three datastes, and seperately on the three datasets) by R and Python, run in 200 iterations. Seurat V3, scMC and LIGER are run in R by the downloaded R packages. Unless specified, all methods are implemented on one 3.4GHz Intel Xeon Gold CPU.

**Table S.12:** Summary of the computation time by scAMACE and other clustering methods for Application 2. The unit of measurement is second. We implemented scAMACE (jointly on the three datastes, and seperately on the three datasets) by R and Python, run in 200 iterations. Seurat V3, scMC and LIGER are run in R by the downloaded R packages. Unless specified, all methods are implemented on one 3.4GHz Intel Xeon Gold CPU.

|  |  |
| --- | --- |
| scAMACE (joint) by R<br>2317.787 | scAMACE (separate) by R<br>249.891(sci-ATAC-Seq)+950.388(scRNA-Seq)+171.823(sc-methyl)=1372.102 |
| scAMACE (joint) by Python<br>418.858 | scAMACE (separate) by Python<br>51.878(sci-ATAC-Seq)+217.728(scRNA-Seq)+25.994(sc-methyl)=295.6 |
| scAMACE (joint) by Python (using GPU)<br>69.652 | scAMACE (separate) by Python (using GPU)<br>7.651(sci-ATAC-Seq)+33.435(scRNA-Seq)+4.161(sc-methyl)=45.247 |
| Seurat V3 (sci-ATAC-Seq+scRNA-Seq)<br>116.688 | scMC(sci-ATAC-Seq+scRNA-Seq+sc-methyl)<br>372.323 |
| LIGER(scRNA-Seq+sc-methyl)<br>1618.965 | LIGER(sci-ATAC-Seq+scRNA-Seq+sc-methyl)<br>80.389 |

**Table S.13:** Summary of the computation time by scAMACE and other clustering methods using 30,000 bootstrap samples (*n*_*acc*_ = *n*_*rna*_ = *n*_*met*_ = 10, 000) from Application 2. The unit of measurement is second. We implemented scAMACE (jointly on the three datastes, and seperately on the three datasets) by R and Python, run in 200 iterations. Seurat V3, scMC and LIGER are run in R by the downloaded R packages. Unless specified, all methods are implemented on one 3.4GHz Intel Xeon Gold CPU.

|  |  |
| --- | --- |
| scAMACE (joint) by R<br>7663.651 | scAMACE (separate) by R<br>1366.109(sci-ATAC-Seq)+2570.669(scRNA-Seq)+1391.464(sc-methyl)=5328.242 |
| scAMACE (joint) by Python<br>1534.631 | scAMACE (separate) by Python<br>307.066(sci-ATAC-Seq)+548.065(scRNA-Seq)+373.092(sc-methyl)=1228.223 |
| scAMACE (joint) by Python (using GPU)<br>250.089 | scAMACE (separate) by Python (using GPU)<br>64.549(sci-ATAC-Seq)+82.53(scRNA-Seq)+49.898(sc-methyl)=196.977 |
| Seurat V3 (sci-ATAC-Seq+scRNA-Seq)<br>290.640 | scMC(sci-ATAC-Seq+scRNA-Seq+sc-methyl)<br>3667.878 |
| LIGER(scRNA-Seq+sc-methyl)<br>5319.259 | LIGER(sci-ATAC-Seq+scRNA-Seq+sc-methyl)<br>555.574 |

